# Convergent Antibody Responses to SARS-CoV-2 Infection in Convalescent Individuals

**DOI:** 10.1101/2020.05.13.092619

**Authors:** Davide F. Robbiani, Christian Gaebler, Frauke Muecksch, Julio C. C. Lorenzi, Zijun Wang, Alice Cho, Marianna Agudelo, Christopher O. Barnes, Anna Gazumyan, Shlomo Finkin, Thomas Hagglof, Thiago Y. Oliveira, Charlotte Viant, Arlene Hurley, Hans-Heinrich Hoffmann, Katrina G. Millard, Rhonda G. Kost, Melissa Cipolla, Kristie Gordon, Filippo Bianchini, Spencer T. Chen, Victor Ramos, Roshni Patel, Juan Dizon, Irina Shimeliovich, Pilar Mendoza, Harald Hartweger, Lilian Nogueira, Maggi Pack, Jill Horowitz, Fabian Schmidt, Yiska Weisblum, Eleftherios Michailidis, Alison W. Ashbrook, Eric Waltari, John E. Pak, Kathryn E. Huey-Tubman, Nicholas Koranda, Pauline R. Hoffman, Anthony P. West, Charles M. Rice, Theodora Hatziioannou, Pamela J. Bjorkman, Paul D. Bieniasz, Marina Caskey, Michel C. Nussenzweig

**Author notes:** Equal contribution (Davide F. Robbiani, Christian Gaebler, Frauke Muecksch, Julio C. C. Lorenzi, Zijun Wang, Alice Cho, Marianna Agudelo, Christopher O. Barnes, Anna Gazumyan, Shlomo Finkin, Thomas Hagglof, Thiago Y. Oliveira, Charlotte Viant). Send correspondence to Paul D. Bieniasz, Marina Caskey, Michel C. Nussenzweig, or Davide F. Robbiani.

## Abstract

During the COVID-19 pandemic, SARS-CoV-2 infected millions of people and claimed hundreds of thousands of lives. Virus entry into cells depends on the receptor binding domain (RBD) of the SARS-CoV-2 spike protein (S). Although there is no vaccine, it is likely that antibodies will be essential for protection. However, little is known about the human antibody response to SARS-CoV-2^1–5^. Here we report on 149 COVID-19 convalescent individuals. Plasmas collected an average of 39 days after the onset of symptoms had variable half-maximal neutralizing titers ranging from undetectable in 33% to below 1:1000 in 79%, while only 1% showed titers >1:5000. Antibody cloning revealed expanded clones of RBD-specific memory B cells expressing closely related antibodies in different individuals. Despite low plasma titers, antibodies to three distinct epitopes on RBD neutralized at half-maximal inhibitory concentrations (IC_50_s) as low as single digit ng/mL. Thus, most convalescent plasmas obtained from individuals who recover from COVID-19 do not contain high levels of neutralizing activity. Nevertheless, rare but recurring RBD-specific antibodies with potent antiviral activity were found in all individuals tested, suggesting that a vaccine designed to elicit such antibodies could be broadly effective.

---

Between April 1 and May 8, 2020, 157 eligible participants enrolled in the study. Of these, 111 (70.7%) were individuals diagnosed with SARS-CoV-2 infection by RT-PCR (cases), and 46 (29.3%) were close contacts of individuals diagnosed with SARS-CoV-2 infection (contacts). While inclusion criteria allowed for enrollment of asymptomatic participants, 8 contacts that did not develop symptoms were excluded from further analyses. The 149 cases and contacts were free of symptoms suggestive of COVID-19 for at least 14 days at the time of sample collection. Participant demographics and clinical characteristics are shown in Table 1 and Extended Data Tables 1 and 2. Only one individual who tested positive for SARS-CoV-2 infection by RT-PCR remained asymptomatic. The other 148 participants reported symptoms suggestive of COVID-19 with an average onset of approximately 39 days (range 17 to 67 days) before sample collection. In this cohort, symptoms lasted for an average of 12 days (0-35 days), and 11 (7%) of the participants were hospitalized. The most common symptoms were fever (83.9%), fatigue (71.1%), cough (62.4%) and myalgia (61.7%) while baseline comorbidities were infrequent (10.7%) (Table 1 and Extended Data Tables 1 and 2). There were no significant differences in duration or severity (see Methods) of symptoms, or in time from onset of symptoms to sample collection between genders or between cases and contacts. There was no age difference between females and males in our cohort (Extended Data Fig. 1).

**Table 1.** Cohort characteristics

| Gender | n | Average age | Case/Contact | Average duration |  | Average Sx Severity (0-10) | ELISA binding (AUC) |  |  |  | Neutralization (NT50) |
| --- | --- | --- | --- | --- | --- | --- | --- | --- | --- | --- | --- |
|  |  |  |  | Sx total | Sx onset to visit |  | RBD |  | S |  |  |
|  |  |  |  |  |  |  | IgG | IgM | IgG | IgM |  |
| Male | 83 | 45<br>(19-76) | 65/18 | 12<br>(0-31) | 39<br>(21-63) | 5.8<br>(0-10) | 2.44 | 1.61 | 4.65 | 1.62 | 867 |
| Female | 66 | 42<br>(19-75) | 46/20 | 12<br>(1-35) | 38<br>(17-67) | 5.4<br>(1-9) | 1.99 | 1.58 | 4.36 | 1.86 | 522 |
| Sx=symptoms |  |  |  |  |  |  |  |  |  |  |  |

Plasma samples were tested for binding to the SARS-CoV-2 RBD and trimeric spike (S) proteins by ELISA using anti-IgG or -IgM secondary antibodies for detection (Fig. 1, Extended Data Table 1 and Extended Data Figs. 2 and 3). Eight independent negative controls and the plasma sample from participant 21 (COV21) were included for normalization of the area under the curve (AUC). Overall, 78% and 70% of the plasma samples tested showed anti-RBD and anti-S IgG AUCs that were at least 2 standard deviations above the control (Fig. 1 a, b). In contrast, only 15% and 34% of the plasma samples showed IgM responses to anti-RBD and anti-S that were at least 2 standard deviations above control, respectively (Fig. 1 c, d). There was no positive correlation between anti-RBD or -S IgG or IgM levels and duration of symptoms or the timing of sample collection relative to onset of symptoms (Fig. 1e, and Extended Data Figs. 3 a-c and 3 g-j). On the contrary, as might be expected, anti-RBD IgM titers were negatively correlated with duration of symptoms and the timing of sample collection (Fig. 1e and Extended Data Fig. 3h). Anti-RBD IgG levels were modestly correlated to age, and the severity of symptoms including hospitalization (Fig. 1 f, g and Extended Data Fig. 3k). Interestingly, females had lower anti-RBD and -S IgG titers than males (Fig. 1h, Extended Data Fig. 2f).

**Figure 1.**
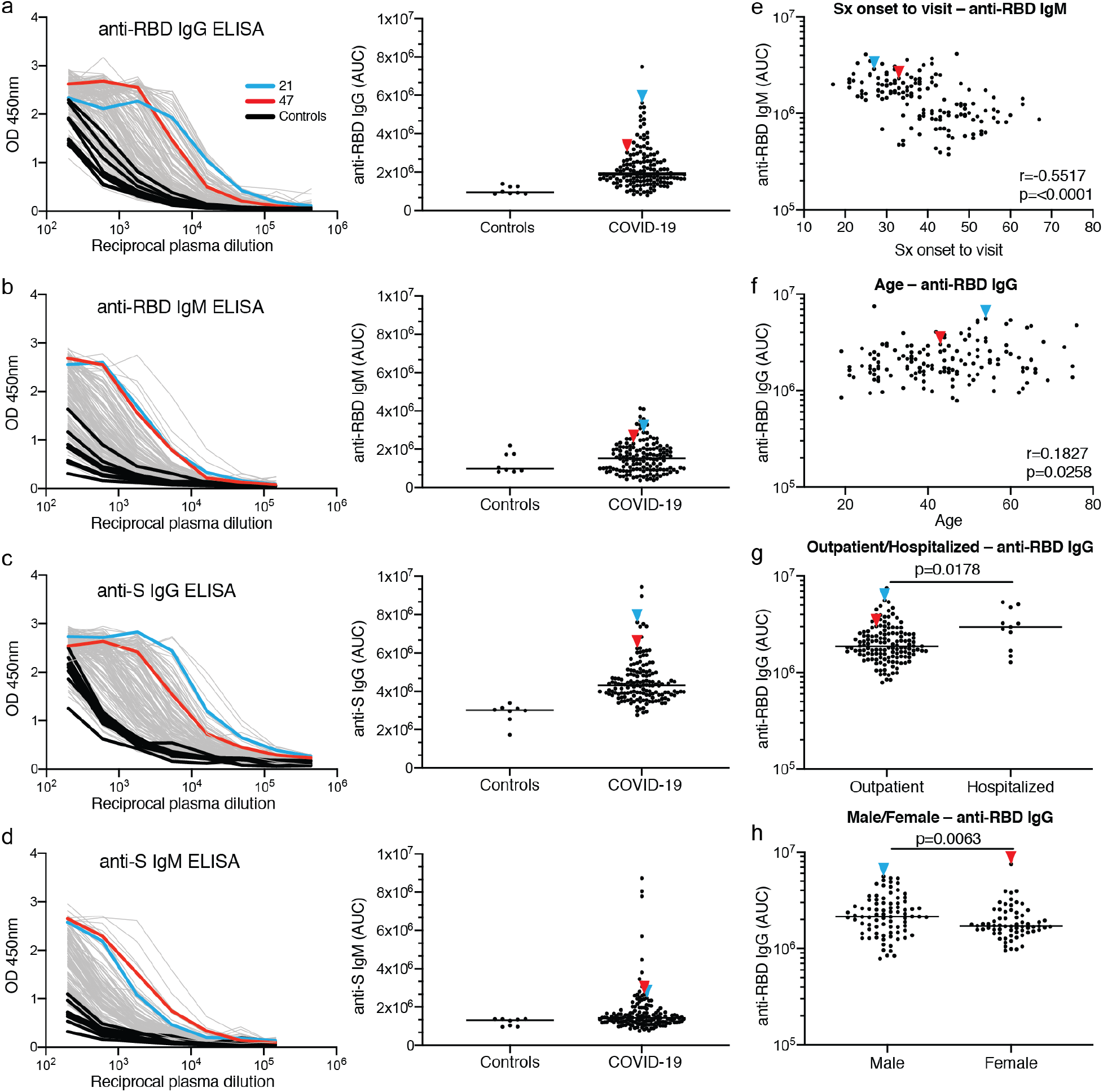
Plasma antibodies against SARS-CoV-2. **a-d,** Graphs show results of ELISAs measuring plasma reactivity to RBD (**a, b**) and S protein (**c, d**). Left shows optical density units at 450 nm (OD, Y axis) and reciprocal plasma dilutions (X axis). Negative controls in black; individuals 21, and 47 in blue and red lines and arrowheads, respectively. Right shows normalized area under the curve (AUC) for controls and each of 149 individuals in the cohort. **e**, Symptom (Sx) onset to time of sample collection in days (X axis) plotted against normalized AUC for IgM binding to RBD (Y axis) r=0.5517 and p=<0.0001. **f**, Participant age in years (X axis) plotted against normalized AUC for IgG binding to RBD (Y axis) r=0.1827 and p=0.0258. The r and p values for the correlations in **e** and **f** were determined by two-tailed Spearman’s. **g,** IgG anti-RBD normalized AUC for outpatients and hospitalized individuals p=0.0178. **h,** IgG anti-RBD normalized AUC for males and females p=0.0063. For **g** and **h** horizontal bars indicate median values. Statistical significance was determined using two-tailed Mann-Whitney U test.

To measure the neutralizing activity in convalescent plasmas we used HIV-1-based virions carrying a nanoluc luciferase reporter that were pseudotyped with the SARS-CoV-2 spike (SARS-CoV-2 pseudovirus, see Methods, Fig. 2 and Extended Data Fig. 4). The overall level of neutralizing activity in the cohort, as measured by the half-maximal neutralizing titer (NT_50_) was generally low, with 33% undetectable and 79% below 1,000 (Fig. 2 a, b). The geometric mean NT_50_ was 121 (arithmetic mean = 714), and only 2 individuals reached NT_50_s above 5,000 (Fig. 2 a, b and Extended Data Table 1).

**Figure 2.**
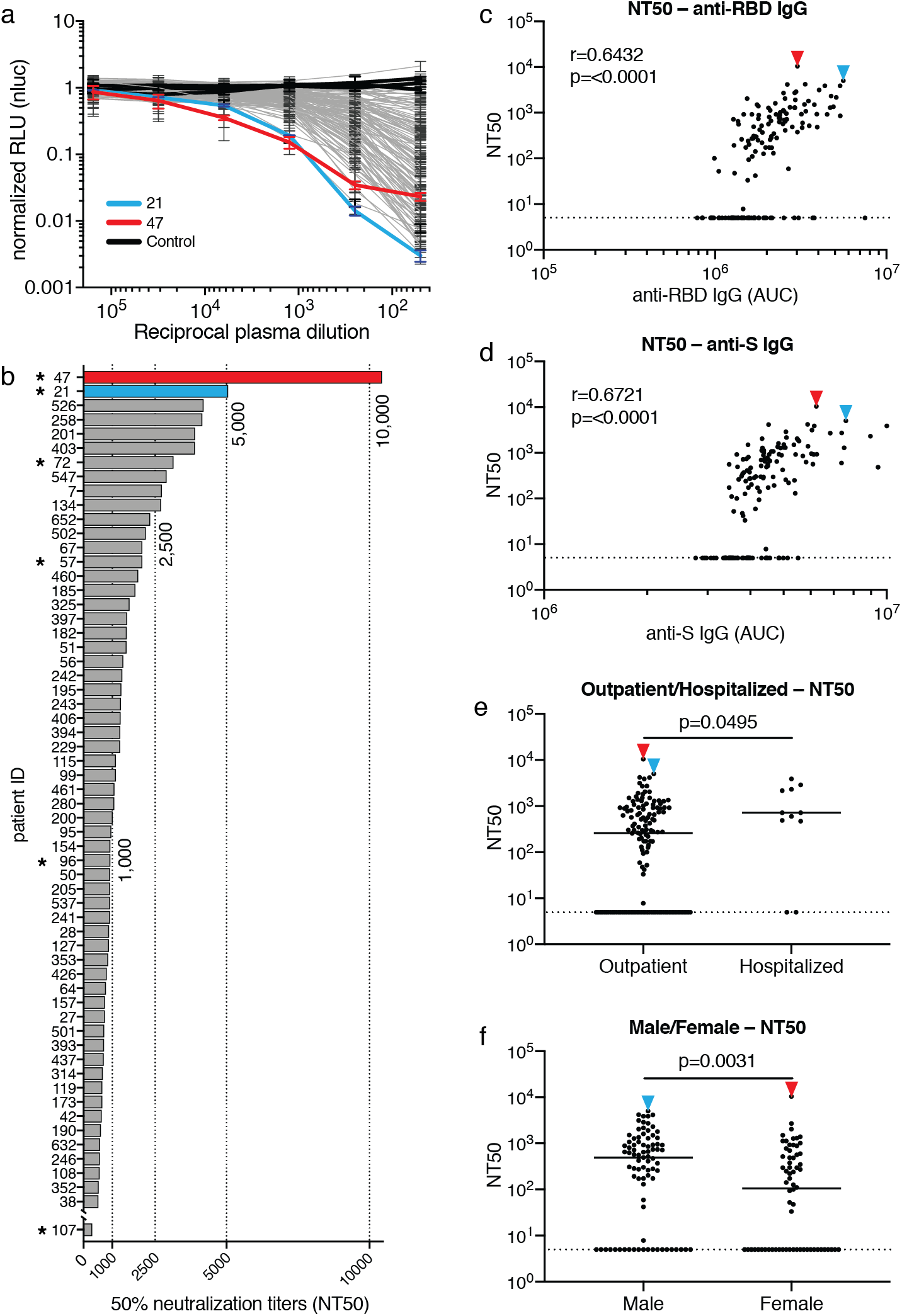
Neutralization of SARS-CoV-2 pseudovirus by plasma. **a,** Graph shows normalized relative luminescence values (RLU, Y axis) in cell lysates of 293T_ACE2_ cells 48 hours after infection with nanoluc-expressing SARS-CoV-2 pseudovirus in the presence of increasing concentrations of plasma (X axis) derived from 149 participants (grey, except individuals 47 and 21 in red, and blue lines, bars and arrowheads, respectively) and 3 negative controls (black lines). Standard deviations of duplicates of one representative experiment are shown. **b**, Ranked average half-maximal inhibitory plasma neutralizing titer (NT_50_) for the 59 of 149 individuals with NT_50_s >500 and individual 107. See also Extended Data Table 1. Asterisks indicate donors from which antibody sequences were derived. **c**, AUC for anti-RBD IgG ELISA (X axis) plotted against NT_50_ (Y axis) r=0.6432, p=<0.0001. **d**, AUC for anti-S IgG ELISA (X axis) plotted against NT_50_ (Y axis) r=0.6721, p=<0.0001. **e**, NT_50_ for outpatients and hospitalized individuals p=0.0495. **f**, NT_50_ for all males and females in the cohort p=0.0031. Dotted line in c to f (NT_50_=5) represents lower limit of detection (LLOD). Samples with undetectable neutralizing titers were plotted at LLOD. Correlations in **c** and **d** were determined by two-tailed Spearman’s. Statistical significance in **e** and **f** was determined using two-tailed Mann-Whitney U test. Horizontal bars indicate median values.

Notably, levels of anti-RBD- and -S IgG antibodies correlated strongly with NT_50_ (Fig. 2 c, d). Neutralizing activity also correlated with age, duration of symptoms and symptom severity (Extended Data Fig. 5). Consistent with this observation, hospitalized individuals with longer symptom duration showed slightly higher average levels of neutralizing activity than non-hospitalized individuals (p=0.0495, Fig. 2e). Finally, we observed a significant difference in neutralizing activity between males and females (p=0.0031, Fig. 2f). The difference between males and females was consistent with higher anti-RBD and -S IgG titers in males, and could not be attributed to age, severity, timing of sample collection relative to onset of symptoms or duration of symptoms (Fig. 1h, Extended Data Fig. 1 a-d and 2f).

To determine the nature of the antibodies elicited by SARS-CoV-2 infection we used flow cytometry to isolate individual B lymphocytes with receptors that bound to RBD from the blood of 6 selected individuals including the 2 top and 4 high to intermediate neutralizers (Fig. 3). The frequency of antigen-specific B cells, identified by their ability to bind to both Phycoerythrin (PE)- and AF647-labeled RBD, ranged from 0.07 to 0.005% of all circulating B cells in COVID-19 convalescents but they were undetectable in pre-COVID-19 controls (Fig. 3a and Extended Data Fig. 6). We obtained 534 paired IgG heavy and light chain (IGH and IGL) sequences by reverse transcription and subsequent PCR from individual RBD-binding B cells from the 6 convalescent individuals (see Methods and Extended Data Table 3). When compared to the human antibody repertoire, several IGHV and IGLV genes were significantly over-represented (Extended Data Fig. 7). The average number of V genes nucleotide mutations for IGH and IGL was 4.2 and 2.8, respectively (Extended Data Fig. 8), which is lower than in antibodies cloned from individuals suffering from chronic infections such as Hepatitis B or HIV-1, and similar to antibodies derived from primary malaria infection or non-antigen-enriched circulating IgG memory cells^6–8^ (Wang et al, in press, https://www.biorxiv.org/content/10.1101/2020.03.04.976159v1.full). Among other antibody features, IGH CDR3 length was indistinguishable from the reported norm and hydrophobicity was below average (Extended Data Fig. 8)^9^.

**Figure 3.**
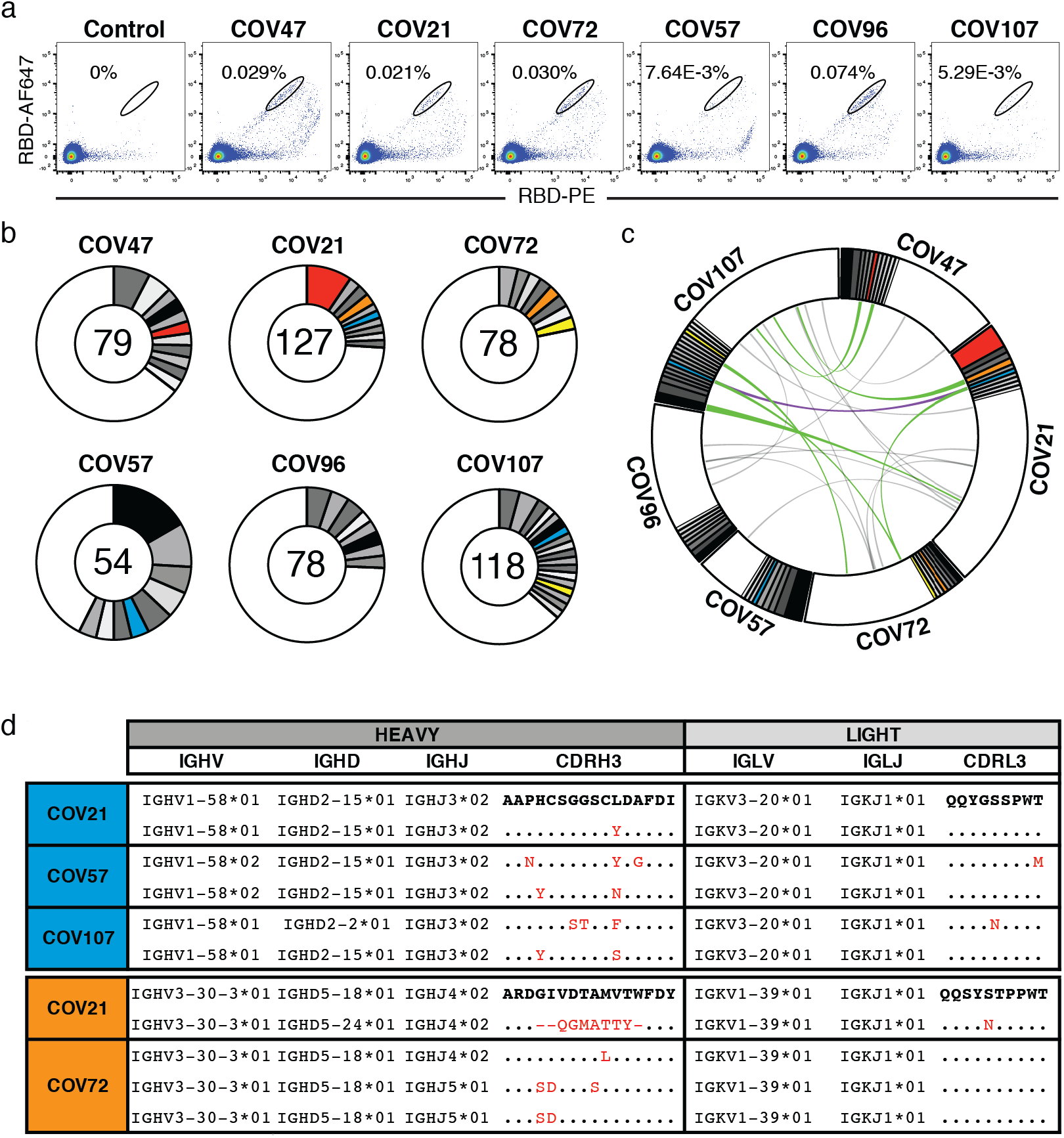
Anti-SARS-CoV-2 RBD antibodies. **a**. Representative flow cytometry plots showing dual AF647- and PE-RBD binding B cells in control and 6 study individuals (for gating strategy see Extended Data Fig. 6). Percentages of antigen specific B cells are indicated. Control is a healthy control sample obtained pre-COVID-19. **b**, Pie charts depicting the distribution of antibody sequences from 6 individuals. The number in the inner circle indicates the number of sequences analyzed for the individual denoted above the circle. White indicates sequences isolated only once, and grey or colored pie slices are proportional to the number of clonally related sequences. Red, blue, orange and yellow pie slices indicate clones that share the same IGHV and IGLV genes. **c**, Circos plot shows sequences from all 6 individuals with clonal relationships depicted as in b. Interconnecting lines indicate the relationship between antibodies that share V and J gene segment sequences at both IGH and IGL. Purple, green and gray lines connect related clones, clones and singles, and singles to each other, respectively. **d**, Sample sequence alignment for antibodies originating from different individuals that display highly similar IGH V(D)J and IGL VJ sequences including CDR3s. Amino acid differences in CDR3s to the bolded reference sequence above are indicated in red and dots represent identities.

As is the case with other human pathogens, there were expanded clones of viral antigen binding B cells in all COVID-19 individuals tested (see Methods and Fig. 3 b,c). Overall, 32.2% of the recovered IGH and IGL sequences were from clonally expanded B cells (range 21.8-57.4% across individuals, Fig. 3b). Antibodies that shared specific combinations of IGHV and IGLV genes in different individuals comprised 14% of all the clonal sequences (colored pie slices in Fig. 3 b,c). Remarkably, the amino acid sequences of some antibodies found in different individuals were nearly identical (Fig. 3 e,d). For example, antibodies expressed by clonally expanded B cells with IGHV1-58/IGKV3-20 and IGHV3-30-3/IGKV1-39 found repeatedly in different individuals had amino acid sequence identities of up to 99% and 92%, respectively (Fig. 3d and Extended Data Table 4). We conclude that the IgG memory response to the SARS-CoV-2 RBD is rich in recurrent and clonally expanded antibody sequences.

To examine the binding properties of anti-SARS-CoV-2 antibodies, we expressed 84 representative antibodies, 56 from clones and 28 from singlets (Extended Data Table 5). ELISA assays showed that 94% (79 out of 84) of the antibodies tested including clonal and unique sequences bound to the SARS-CoV-2 RBD with an average half-maximal effective concentration (EC_50_) of 6.1 ng/mL (Fig. 4a and Extended Data Fig. 9a). A fraction of those (7 out of 79, or 9%) cross-reacted with the RBD of SARS-CoV with a mean EC_50_ of 120.1 ng/mL (Extended Data Fig. 9b and c). No significant cross-reactivity was noted to the RBDs of MERS, HCoV-OC43, HCoV-229E or HCoV-NL63.

**Figure 4.**
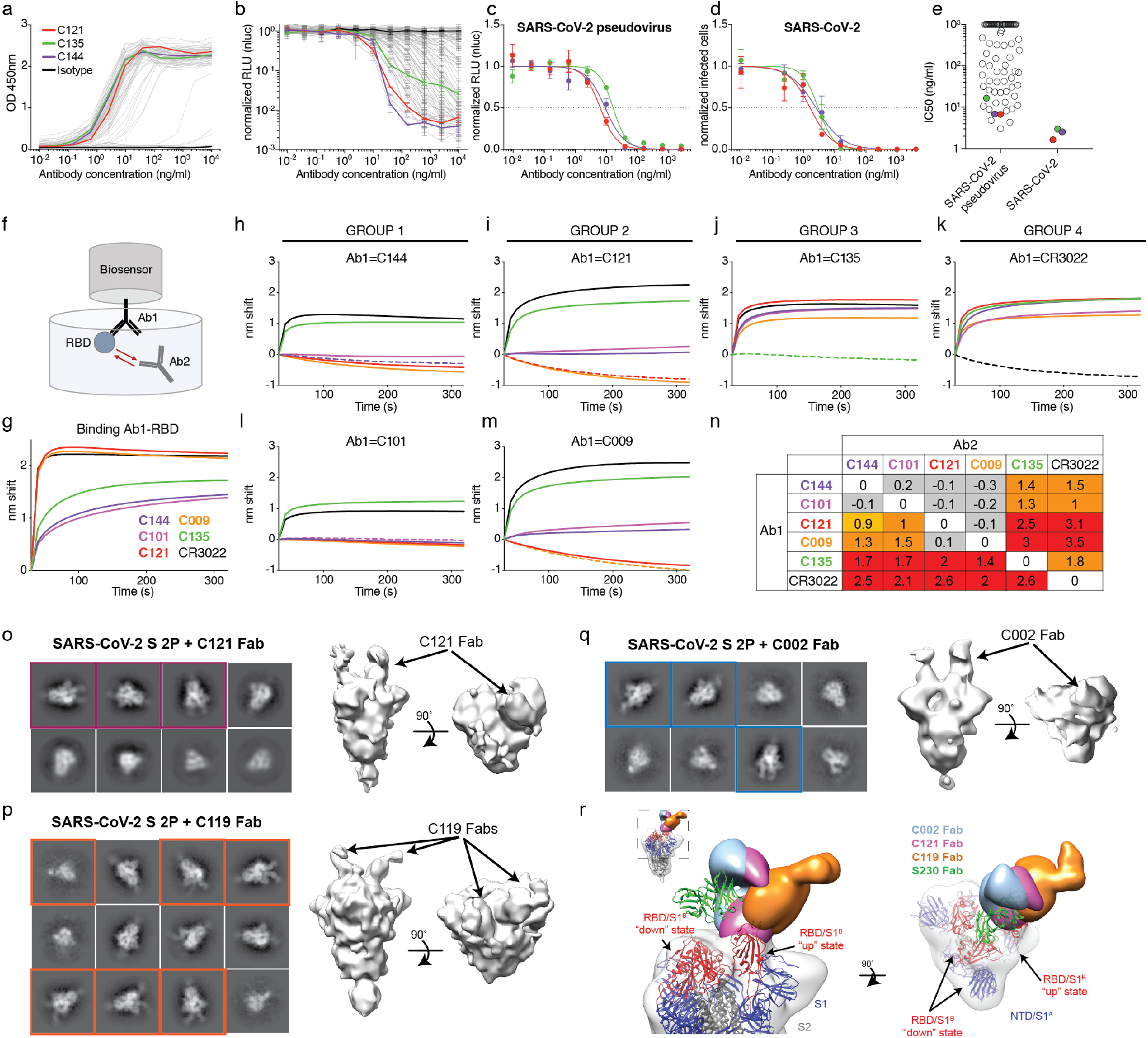
Anti-SARS-CoV-2 RBD antibody reactivity. **a**, Graph show results of ELISA assays measuring monoclonal antibody reactivity to RBD. Optical density units at 450 nm (OD, Y axis) vs. antibody concentrations (X axis). C121, C135 C144 and isotype control in red, green, purple, and black respectively, in all panels. **b**, Graph shows normalized relative luminescence values (RLU, Y axis) in cell lysates of 293T_ACE2_ cells 48 hours after infection with SARS-CoV-2 pseudovirus in the presence of increasing concentrations of monoclonal antibodies (X axis). **c**, RLU for SARS-CoV-2 pseudovirus assay (Y axis) vs. titration of monoclonal antibodies C121, C135 and C144 in one of two independent experiments (see Extended Data Table 6). **d**, SARS-CoV-2 real virus neutralization assay. Infected cells (Y axis) vs. titration of monoclonal antibodies C121, C135 and C144 in two independent experiments. For a and b panels, isotype control antibody in black. e, IC_50_s for antibodies assayed in **b** and **d. f**, Diagrammatic representation of biolayer interferometry experiment. **g**, Graph shows binding of C144, C101, C121, C009, C135, and CR3022^10,11^ to RBD. **h-m**, Secondary antibody binding to preformed IgG-RBD complexes (Ab1). The table displays the shift in nanometers after second antibody (Ab2) binding to the antigen in the presence of the first antibody (Ab1). Values are normalized by the subtraction of the autologous antibody control. **o-q**, Representative 2D-class averages and 3D reconstructed volumes for SARS-CoV-S 2P trimers complexed with C002, C119, and C121 Fabs. 2D-class averages with observable Fab density are boxed. **r**, Overlay of S-Fab complexes with fully-occupied C002 (blue), C121 (magenta) and C119 (orange) Fabs aligned on the RBD “up” conformational state. The SARS-CoV-2 S model with 1 “up” RBD state (PDB 6VYB) was fit into the density and the SARS-CoV antibody S230 (PDB 6NB6) shown as reference (green ribbon).

To determine whether the monoclonal antibodies have neutralizing activity, we tested them against the SARS-CoV-2 pseudovirus (Fig. 4 and Extended Data Table 6). Among 79 RBD binding antibodies tested, we found 40 that neutralized SARS-CoV-2 pseudovirus with nanogram per milliliter half-maximal inhibitory concentrations (IC_50_s) ranging from 3 to 709 (Fig. 4 b,c and e, Extended Data Table 6). A subset of the most potent of these antibodies were also tested against authentic SARS-CoV-2 and neutralized with IC_50_s of less than 5 ng/ml (Fig. 4 d,e).

Potent neutralizing antibodies were found in individuals irrespective of their plasma NT_50_s. For example, C121, C144, and C135 with IC_50_s of 1.64, 2.55 and 2.98 ng/mL against authentic SARS-CoV-2, respectively, were obtained from individuals COV107, COV47, and COV72 whose plasma NT_50_ values were of 297, 10,433 and 3,138, respectively (Figs. 2b and 4). Finally, clones of antibodies with shared IGHV and IGLV genes were among the best neutralizers, e.g., antibody C002 composed of IGHV3-30/IGKV1-39 is shared by the 2 donors with the best plasma neutralizing activity (red pie slice in Fig. 3b and Fig. 4). We conclude that even individuals with modest plasma neutralizing activity harbor rare IgG memory B cells that produce potent SARS-CoV-2 neutralizing antibodies.

To determine whether human anti-SARS-CoV-2 monoclonal antibodies with neutralizing activity can bind to distinct domains on the RBD, we performed bilayer interferometry experiments in which a preformed antibody-RBD immune complex was exposed to a second monoclonal. The antibodies tested comprised 3 groups, all of which differ in their binding properties from CR3022, an antibody that neutralizes SARS-CoV and binds to, but does not neutralize SARS-CoV-2^10,11^. Representatives of each of the 3 groups include: C144 and C101 in Group 1; C121 and C009 in Group 2; C135 in Group 3. All of these antibodies can bind after CR3022. Groups 1 and 2 also bind after Group 3, and Groups 1 and 2 differ in that Group 1 can bind after Group 2 but not vice versa (Fig. 4 f-n). We conclude that similar to SARS-CoV, there are multiple distinct neutralizing epitopes on the RBD of SARS-CoV-2.

To further define the binding characteristics of Groups 1 and 2 antibodies, we imaged SARS-CoV-2 S–Fab complexes by negative stain electron microscopy (nsEM) using C002 (Group 1, an IGHV3-30/IGKV1-39 antibody, which is clonally expanded in 2 donors), C119 and C121 (both in Group 2) Fabs (Fig. 4 f-r and Extended Data Fig. 10). Consistent with the conformational flexibility of the RBD, 2D class averages showed heterogeneity in both occupancy and conformation of bound Fabs for both groups (Fig. 4o-q). The 3D reconstructions of S-Fab complexes revealed both a fully-occupied RBD and partial density for a second bound Fab, suggesting that Fabs from both groups are able to recognize “up” and “down” states of the RBD as previously described for some of the antibodies targeting this epitope^12,13^. The 3D reconstructions are also consistent with BLI measurements indicating that Groups 1 and 2 antibodies bind a RBD epitope distinct from antibody CR3022, which is only accessible on the RBD “up” conformational state^11^, and bind the RBD with different angles of approach, with Group 1 antibodies most similar to the approach angle of the SARS-CoV antibody S230 (Fig. 4r)^14^.

Human monoclonal antibodies with neutralizing activity against pathogens ranging from viruses to parasites have been obtained from naturally infected individuals by single cell antibody cloning. Several have been shown to be effective in protection and therapy in model organisms and in early phase clinical studies, but only one antiviral monoclonal is currently in clinical use^15^. Antibodies are relatively expensive and more difficult to produce than small molecule drugs. However, they differ from drugs in that they can engage the host immune system through their constant domains that bind to Fc gamma receptors on host immune cells^16^. These interactions can enhance immunity and help clear the pathogen or infected cells, but they can also lead to disease enhancement during Dengue^17^ and possibly coronavirus infections^18^. This problem has impeded Dengue vaccine development but would not interfere with the clinical use of potent neutralizing antibodies that can be modified to prevent Fc gamma receptor interactions and remain protective against viral pathogens^19^.

Antibodies are essential elements of most vaccines and will likely be crucial component of an effective vaccine against SARS-CoV-2^20^(PMID:32434945; PMID:32434946). Recurrent antibodies have been observed in other infectious diseases and vaccinal responses ^21–24^(Wang et al, in press, https://www.biorxiv.org/content/10.1101/2020.03.04.976159v1.full). The observation that plasma neutralizing activity is low in most convalescent individuals, but that recurrent anti-SARS-CoV-2 RBD antibodies with potent neutralizing activity can be found in individuals with unexceptional plasma neutralizing activity suggests that humans are intrinsically capable of generating anti-RBD antibodies that potently neutralize SARS-CoV-2. Thus, vaccines that selectively and efficiently induce antibodies targeting the SARS-CoV-2 RBD may be especially effective.

## Methods

### Study participants

Study participants were recruited at the Rockefeller University Hospital in New York from April 1 through May 8, 2020. Eligible participants were adults aged 18-76 years who were either diagnosed with SARS-CoV-2 infection by RT-PCR and were free of symptoms of COVID-19 for at least 14 days (cases), or who were close contacts (e.g., household, co-workers, members of same religious community) with someone who had been diagnosed with SARS-CoV-2 infection by RT-PCR and were free of symptoms suggestive of COVID-19 for at least 14 days (contacts). Exclusion criteria included presence of symptoms suggestive of active SARS-CoV-2 infection, or hemoglobin < 12 g/dL for males and < 11 g/dL for females.

Most study participants were residents of the Greater New York City tri-state region and were enrolled sequentially according to eligibility criteria. Participants were first interviewed by phone to collect information on their clinical presentation, and subsequently presented to the Rockefeller University Hospital for a single blood sample collection. Participants were asked to rate the highest severity of their symptoms on a numeric rating scale ranging from 0 to 10. The score was adapted from the pain scale chart, where 0 was the lack of symptoms, 4 were distressing symptoms (e.g. fatigue, myalgia, fever, cough, shortness of breath) that interfered with daily living activities, 7 were disabling symptoms that prevented the performance of daily living activities, and 10 was unimaginable/unspeakable discomfort (in this case, distress due to shortness of breath). All participants provided written informed consent before participation in the study and the study was conducted in accordance with Good Clinical Practice.

### Blood samples processing and storage

Peripheral Blood Mononuclear Cells (PBMCs) were obtained by gradient centrifugation and stored in liquid nitrogen in the presence of FCS and DMSO. Heparinized plasma and serum samples were aliquoted and stored at −20°C or less. Prior to experiments, aliquots of plasma samples were heat-inactivated (56C for 1 hour) and then stored at 4C.

### Cloning, expression and purification of recombinant coronavirus proteins

Codon-optimized nucleotide sequences encoding the SARS-CoV-2 S ectodomain (residues 16-1206) and receptor binding domain (RBD; residues 331-524) were synthesized and subcloned into the mammalian expression pTwist-CMV BetaGlobin vector by Twist Bioscience Technologies based on an early SARS-CoV-2 sequence isolate (GenBank MN985325.1). The SARS-CoV-2 RBD construct included an N-terminal human IL-2 signal peptide and dual C-terminal tags ((GGGGS)2-HHHHHHHH (octa-histidine), and GLNDIFEAQKIEWHE (AviTag)). In addition, the corresponding S1^B^ or receptor binding domains for SARS-CoV S (residues 318-510; GenBank AAP13441.1), MERS-CoV S (residues 367-588; GenBank JX869059.2), HCoV-NL63 (residues 481-614; GenBank AAS58177.1), HCoV-OC43 (residues 324-632; GenBank AAT84362.1), and HCoV-229E (residues 286-434; GenBank AAK32191.1) were synthesized with the same N-and C-terminal extensions as the SARS-CoV-2 RBD construct and subcloned into the mammalian expression pTwist-CMV BetaGlobin vector (Twist Bioscience Technologies). The SARS-CoV-2 S ectodomain was modified as previously described^4^ Briefly, the S ectodomain construct included an N-terminal mu-phosphatase signal peptide, 2P stabilizing mutations (K986P and V987P), mutations to remove the S1/S2 furin cleavage site (_682_RRAR_685_ to GSAS), a C-terminal extension (IKGSG-RENLYFQG (TEV protease site), GGGSG-YIPEAPRDGQAYVRKDGEWVLLSTFL (foldon trimerization motif), G-HHHHHHHH (octa-histidine tag), and GLNDIFEAQKIEWHE (AviTag)). The SARS-CoV-2 S 2P ectodomain and RBD constructs were produced by transient transfection of 500 mL of Expi293 cells (Thermo Fisher) and purified from clarified transfected cell supernatants four days post-transfection using Ni^2+^-NTA affinity chromatography (GE Life Sciences). Affinity-purified proteins were concentrated and further purified by size-exclusion chromatography (SEC) using a Superdex200 16/60 column (GE Life Sciences) running in 1x TBS (20 mM Tris-HCl pH 8.0, 150 mM NaCl, and 0.02% NaN_3_). Peak fractions were analyzed by SDS-PAGE, and fractions corresponding to soluble S 2P trimers or monomeric RBD proteins were pooled and stored at 4°C.

### ELISAs

ELISAs to evaluate antibodies binding to SARS-CoV-2 RBD and trimeric spike proteins were performed by coating of high binding 96 half well plates (Corning #3690) with 50 μL per well of a 1μg/mL protein solution in PBS overnight at 4°C. Plates were washed 6 times with washing buffer (1xPBS with 0.05% Tween 20 (Sigma-Aldrich)) and incubated with 170 μL per well blocking buffer (1xPBS with 2% BSA and 0.05% Tween20 (Sigma)) for 1 hour at room temperature (RT). Immediately after blocking, monoclonal antibodies or plasma samples were added in PBS and incubated for 1 hr at RT. Plasma samples were assayed at a 1:200 starting dilution and seven additional 3-fold serial dilutions. Monoclonal antibodies were tested at 10μg/ml starting concentration and 10 additional 4-fold serial dilutions. Plates were washed 6 times with washing buffer and then incubated with anti-human IgG or IgM secondary antibody conjugated to horseradish peroxidase (HRP) (Jackson Immuno Research 109-036-088 and 109-035-129) in blocking buffer at a 1:5000 dilution. Plates were developed by addition of the HRP substrate, TMB (ThermoFisher) for 10 minutes, then the developing reaction was stopped by adding 50μl 1M H2SO4 and absorbance was measured at 450nm with an ELISA microplate reader (FluoStar Omega, BMG Labtech). For plasma samples, a positive control (plasma from patient COV21, diluted 200-fold in PBS) was added in duplicate to every assay plate. The average of its signal was used for normalization of all the other values on the same plate with Excel software prior to calculating the area under the curve using Prism 8 (GraphPad). For monoclonal antibodies, the half-maximal effective concentration (EC_50_) was determined using 4-parameter nonlinear regression (GraphPad Prism).

### 293T_ACE2_ cells

For constitutive expression of ACE2 in 293T cells, a cDNA encoding ACE2, carrying two inactivating mutations in the catalytic site (H374N & H378N), was inserted into CSIB 3’ to the SFFV promoter^25^. 293T_ACE2_ cells were generated by transduction with CSIB based virus followed by selection with 5 μg/ml Blasticidin.

### SARS-CoV-2 and SARS-CoV pseudotyped reporter viruses

A plasmid expressing a C-terminally truncated SARS-CoV-2 S protein (pSARS-CoV2-S_trunc_) was generated by insertion of a human-codon optimized cDNA encoding SARS-CoV-2 S lacking the C-terminal 19 codons (Geneart) into pCR3.1. The S ORF was taken from “Wuhan seafood market pneumonia virus isolate Wuhan-Hu-1” (NC_045512). For expression of full-length SARS-CoV S protein, “Human SARS coronavirus Spike glycoprotein Gene ORF cDNA clone expression plasmid (Codon Optimized)” (here referred to as pSARS-CoV-S) was obtained from SinoBiological (Cat: VG40150-G-N). An *env*-inactivated HIV-1 reporter construct (pNL4-3ΔEnv-nanoluc) was generated from pNL4-3^26^ by introducing a 940 bp deletion 3’ to the *vpu* stop-codon, resulting in a frameshift in *env.* The human codon-optimized nanoluc Luciferase reporter gene (*Nluc,* Promega) was inserted in place of nucleotides 1-100 of the *nef-gene.* To generate pseudotyped viral stocks, 293T cells were transfected with pNL4-3ΔEnv-nanoluc and pSARS-CoV2-S_trunc_ or pSARS-CoV-S using polyethyleneimine. Co-transfection of pNL4-3ΔEnv-nanoluc and S-expression plasmids leads to production of HIV-1-based virions carrying either the SARS-CoV-2 or SARS-CoV spike protein on the surface. Eight hours after transfection, cells were washed twice with PBS and fresh media was added. Supernatants containing virions were harvested 48 hours post transfection, filtered and stored at −80°C. Infectivity of virions was determined by titration on 293T_ACE2_ cells.

### Pseudotyped virus neutralization assay

Five-fold serially diluted plasma from COVID-19 convalescent individuals and healthy donors or four-fold serially diluted monoclonal antibodies were incubated with the SARS-CoV-2 or SARS-CoV pseudotyped virus for 1 hour at 37°C degrees. The mixture was subsequently incubated with 293T_ACE2_ cells for 48 hours after which cells were washed twice with PBS and lysed with Luciferase Cell Culture Lysis 5x reagent (Promega). Nanoluc Luciferase activity in lysates was measured using the Nano-Glo Luciferase Assay System (Promega). Relative luminescence units obtained were normalized to those derived from cells infected with SARS-CoV-2 or SARS-CoV pseudotyped virus in the absence of plasma or monoclonal antibodies. The half-maximal inhibitory concentration for plasma (NT_50_) or monoclonal antibodies (IC50) was determined using 4-parameter nonlinear regression (GraphPad Prism).

### Cell lines, virus and virus titration

VeroE6 kidney epithelial cells (*Chlorocebus sabaeus*) and Huh-7.5 hepatoma cells (*H. Sapiens*) were cultured in Dulbecco’s Modified Eagle Medium (DMEM) supplemented with 1% nonessential amino acids (NEAA) and 10% fetal bovine serum (FBS) at 37°C and 5% CO_2_. All cell lines have been tested negative for contamination with mycoplasma and were obtained from the ATCC (with the exception for Huh-7.5).

SARS-CoV-2, strain USA-WA1/2020, was obtained from BEI Resources and amplified in VeroE6 cells at 33°C. Viral titers were measured on Huh-7.5 cells by standard plaque assay (PA). Briefly, 500 μL of serial 10-fold virus dilutions in Opti-MEM were used to infect 400,000 cells seeded the day prior in a 6-well plate format. After 90 min adsorption, the virus inoculum was removed, and cells were overlayed with DMEM containing 10% FBS with 1.2% microcrystalline cellulose (Avicel). Cells were incubated for five days at 33°C, followed by fixation with 3.5% formaldehyde and crystal violet staining for plaque enumeration. All experiments were performed in a biosafety level 3 laboratory.

### Microscopy-based neutralization assay of authentic SARS-CoV-2

The day prior to infection VeroE6 cells were seeded at 12,500 cells/well into 96-well plates. Antibodies were serially diluted in BA-1, mixed with a constant amount of SARS-CoV-2 (grown in VeroE6) and incubated for 60 min at 37°C. The antibody-virus-mix was then directly applied to VeroE6 cells (MOI of ~0.1 PFU/cell). Cells were fixed 18 hours post infection by adding an equal volume of 7% formaldehyde to the wells, followed by permeabilization with 0.1% Triton X-100 for 10 min. After extensive washing, cells were incubated for 1 hour at room temperature with blocking solution of 5% goat serum in PBS (catalog no. 005–000–121; Jackson ImmunoResearch). A rabbit polyclonal anti-SARS-CoV-2 nucleocapsid antibody (catalog no. GTX135357; GeneTex) was added to the cells at 1:500 dilution in blocking solution and incubated at 4°C overnight. A goat anti-rabbit AlexaFluor 594 (catalog no. A-11012; Life Technologies) at a dilution of 1:2,000 was used as a secondary antibody. Nuclei were stained with Hoechst 33342 (catalog no. 62249; Thermo Scientific) at a 1:1,000 dilution. Images were acquired with a fluorescence microscope and analyzed using ImageXpress Micro XLS (Molecular Devices, Sunnyvale, CA). All statistical analyses were done using Prism 8 software (GraphPad).

### Biotinylation of viral protein for use in flow cytometry

Purified and Avi-tagged SARS-CoV-2 RBD was biotinylated using the Biotin-Protein Ligase-BIRA kit according to manufacturer’s instructions (Avidity). Ovalbumin (Sigma, A5503-1G) was biotinylated using the EZ-Link Sulfo-NHS-LC-Biotinylation kit according to the manufacturer’s instructions (Thermo Scientific). Biotinylated Ovalbumin was conjugated to streptavidin-BV711 (BD biosciences, 563262) and RBD to streptavidin-PE (BD biosciences, 554061) and streptavidin-Alexa Fluor 647 (AF647, Biolegend, 405237) respectively^27^.

### Single cell sorting by flow cytometry

PBMCs were enriched for B cells by negative selection using a pan B cell isolation kit according to the manufacturer’s instructions (Miltenyi Biotec, 130-101-638). The enriched B cells were incubated in FACS buffer (1 X Phosphate-buffered Saline (PBS), 2% calf serum, 1 mM EDTA) with the following anti-human antibodies: anti-CD20-PECy7 (BD Biosciences, 335793), anti-CD3-APC-eFluro 780 (Invitrogen, 47-0037-41), anti-CD8-APC-eFluro 780 (Invitrogen, 47-0086-42), anti-CD16-APC-eFluro 780 (Invitrogen, 47-0168-41), anti-CD14-APC-eFluro 780 (Invitrogen, 47-0149-42), as well as Zombie NIR (BioLegend, 423105), and fluorophore-labeled RBD and Ovalbumin for 30 minutes on ice^27^. Single CD3^-^CD8^-^CD16^-^ CD20^+^Ova^-^RBD-PE^+^RBD-AF647^+^ B cells were sorted into individual wells of a 96-well plates containing 4 μl of lysis buffer (0.5 X PBS, 10mM DTT, 3000 units/mL RNasin Ribonuclease Inhibitors (Promega, N2615) per well using a FACS Aria III (Becton Dickinson). The sorted cells were frozen on dry ice, and then stored at −80°C or immediately used for subsequent RNA reverse transcription. Although cells were not stained for IgG expression, they are memory B cells based on the fact that they are CD20+ (a marker absent in plasmablasts) and they express IgG (since antibodies were amplified from these cells using IgG-specific primers).

### Antibody sequencing, cloning and expression

Antibodies were identified and sequenced as described previously^22,28,29^. Briefly, RNA from single cells was reverse-transcribed (SuperScript III Reverse Transcriptase, Invitrogen, 18080-044) and the cDNA stored at −20°C or used for subsequent amplification of the variable IGH, IGL and IGK genes by nested PCR and Sanger sequencing^28^. Amplicons from the first PCR reaction were used as templates for Sequence- and Ligation-Independent Cloning (SLIC) into antibody expression vectors. Recombinant monoclonal antibodies and Fabs were produced and purified as previously described^30,31^.

### Biolayer interferometry

BLI assays were performed on the Octet Red instrument (ForteBio) at 30°C with shaking at 1,000 r.p.m. Epitope binding assays were performed with protein A biosensor (ForteBio 18-5010), following the manufacture protocol “classical sandwich assay”. (1) Sensor check: sensors immersed 30 sec in buffer alone (buffer ForteBio 18-1105). (2) Capture 1^st^ Ab: sensors immersed 10min with Ab1 at 40 μg/mL. (3) Baseline: sensors immersed 30sec in buffer alone. (4) Blocking: sensors immersed 5 min with IgG isotype control at 50 μg/mL. (6) Antigen association: sensors immersed 5 min with RBD at 100 μg/mL. (7) Baseline: sensors immersed 30 sec in buffer alone. (8) Association Ab2: sensors immersed 5min with Ab2 at 40 μg/mL. Curve fitting was performed using the Data analysis software (ForteBio).

### Computational analyses of antibody sequences

Antibody sequences were trimmed based on quality and annotated using Igblastn v1.14.0^32^ with IMGT domain delineation system. Annotation was performed systematically using Change-O toolkit v.0.4.5^33^. Heavy and light chains derived from the same cell were paired, and clonotypes were assigned based on their V and J genes using in-house R and Perl scripts (Fig. 3 b,c). All scripts and the data used to process antibody sequences are publicly available on GitHub (https://github.com/stratust/igpipeline).

The frequency distributions of human V genes in anti-SARS-CoV-2 antibodies from this study was compared to Sequence Read Archive SRP010970^34^. The V(D)J assignments were done using IMGT/High V-Quest and the frequencies of heavy and light chain V genes were calculated for 14 and 13 individuals, respectively, using sequences with unique CDR3s. The two-tailed t test with unequal variances was used to determine statistical significance (Extended Data Fig. 7).

Nucleotide somatic hypermutation and CDR3 length were determined using in-house R and Perl scripts. For somatic hypermutations, IGHV and IGLV nucleotide sequences were aligned against their closest germlines using IgBlast and the number of differences were considered nucleotide mutations. The average mutations for V genes was calculated by dividing the sum of all nucleotide mutations across all patients by the number of sequences used for the analysis. To calculate the GRAVY scores of hydrophobicity^35^ we used Guy H.R. Hydrophobicity scale based on free energy of transfer (kcal/mole)^36^ implemented by the R package Peptides available in the Comprehensive R Archive Network repository (https://journal.r-project.org/archive/2015/RJ-2015-001/RJ-2015-001.pdf). We used 534 heavy chain CDR3 amino acid sequences from this study and 22,654,256 IGH CDR3 sequences from the public database of memory B-cell receptor sequences^37^. The Shapiro-Wilk test was used to determine whether the GRAVY scores are normally distributed. The GRAVY scores from all 533 IGH CDR3 amino acid sequences from this study (sequence COV047_P4_IgG_51-P1369 lacks CDR3 amino acid sequence) were used to perform the test and 5000 GRAVY scores of the sequences from the public database were randomly selected. The Shapiro-Wilk p-values were 6.896 x 10^-3^ and 2.217 x 10^-6^ for sequences from this study and the public database, respectively, indicating the data are not normally distributed. Therefore, we used the Wilcoxon non-parametric test to compare the samples, which indicated a difference in hydrophobicity distribution (p = 5 x 10^-6^; Extended Data Fig. 8).

### Negative-stain EM Data Collection and Processing

Purified Fabs (C002, C119, and C121) were complexed with SARS-CoV-2 S trimer at a 2-fold molar excess for 1 min and diluted to 40 μg/mL in TBS immediately before adding 3 μL to a freshly-glow discharged ultrathin, 400 mesh carbon-coated copper grid (Ted Pella, Inc.). Samples were blotted after a 1 min incubation period and stained with 1% uranyl formate for an additional minute before imaging. Micrographs were recorded on a Thermo Fisher Talos Arctica transmission electron microscope operating at 200 keV using a K3 direct electron detector (Gatan, Inc) and SerialEM automated image acquisition software^38^. Images were acquired at a nominal magnification of 28,000x (1.44 Å/pixel size) and a −1.5 to −2.0 μm defocus range. Images were processed in cryoSPARC v2.14, and reference-free particle picking was completed using a gaussian blob picker^39^. Reference-free 2D class averages and *ab initio* volumes were generated in cryoSPARC, and subsequently 3D-classified to identify classes of S-Fab complexes, that were then homogenously refined. Figures were prepared using UCSF Chimera^40^.

## Acknowledgements

We thank all study participants who devoted time to our research; Drs. Barry Coller and Sarah Schlesinger, the Rockefeller University Hospital Clinical Research Support Office and nursing staff. Dr. Joseph L. DeRisi for facilitating interactions with the Chan Zuckerberg BioHub. All members of the M.C.N. laboratory for helpful discussions, Drs. Amelia Escolano, Gaёlle Breton and Bernardo Reis, and Maëa Jankovic for laboratory support. This work was supported by NIH grant P01-AI138398-S1 (M.C.N., C.M.R., P.J.B.) and 2U19AI111825 (M.C.N. and C.M.R).; the Caltech Merkin Institute for Translational Research and P50 AI150464 (P.J.B.), George Mason University Fast Grant and the European ATAC consortium (EC 101003650) to D.F.R.; 3 R01-AI091707-10S1 to C.M.R.; R37-AI64003 to P.D.B.; R01AI78788 to T.H.; The G. Harold and Leila Y. Mathers Charitable Foundation to C.M.R. Electron microscopy was performed in the Caltech Beckman Institute Resource Center for Transmission Electron Microscpy (Drs. Songye Chen and Andrey Malyutin, Directors). C.G. was supported by the Robert S. Wennett Post-Doctoral Fellowship, in part by the National Center for Advancing Translational Sciences (National Institutes of Health Clinical and Translational Science Award program, grant UL1 TR001866), and by the Shapiro-Silverberg Fund for the Advancement of Translational Research. P.D.B. and M.C.N. are Howard Hughes Medical Institute Investigators.

## Contributions

D.F.R., P.D.B., P.J.B., T.H., C.M.R. and M.C.N. conceived, designed and analyzed the experiments. D.F.R., M.C. and C.G. designed clinical protocols. F.M., J.C.C.L., Z.W., A.C., M.A., C.O.B., S.F., T.H., C.V., K.G., F.B., S.T.C., P.M., H.H., L.N., F.S., Y.W., H.-H.H., E.M., A.W.A., K.E.H.T., N.K. and P.R.H. carried out all experiments. A.G. and M.C. produced antibodies. C.O.B., J.P. and E.W. produced SARS-CoV-2 proteins. A.H., R.K., J.H., K.G.M., C.G. and M.C. recruited participants and executed clinical protocols. R.P., J.D., M.P. and I.S. processed clinical samples. T.Y.O., A.P.W. and V.R. performed bioinformatic analysis. D.F.R., P.D.B., P.J.B., T.H., C.M.R. and M.C.N. wrote the manuscript with input from all co-authors.

## Extended Data Tables

**Extended Data Table 1.** Individual participant dem ographics and clinical characteristics

| ID | Age | Gender | Race | Ethnicity | Case/Contact | Duration (days) |  | Sx<br>Severity<br>(0-10) | ELISA binding (AUC) |  |  |  | Neutralization<br>(NT50) |
| --- | --- | --- | --- | --- | --- | --- | --- | --- | --- | --- | --- | --- | --- |
|  |  |  |  |  |  | Sx<br>total | Sx onset<br>to visit |  | RBD |  | S |  |  |
|  |  |  |  |  |  |  |  |  | IgG | IgM | IgG | IgM |  |
| 5 | 43 | M | White | Non-Hispanic | Case | 9 | 41 | 9 | 2.52 | 2.35 | 5.51 | 2.43 | 5.0 |
| 7 | 40 | M | White | Non-Hispanic | Case | 11 | 30 | 6 | 2.92 | 2.54 | 7.39 | 2.37 | 2730.4 |
| 8 | 37 | M | White | Non-Hispanic | Case | 3 | 57 | 5 | 2.11 | 0.81 | 4.46 | 0.88 | 5.0 |
| 9 | 35 | F | White | Non-Hispanic | Case | 12 | 54 | 5 | 2.90 | 1.07 | 4.44 | 2.80 | 5.0 |
| 12 | 27 | F | White | Non-Hispanic | Contact | 7 | 24 | 3 | 1.47 | 1.71 | 4.31 | 1.23 | 5.0 |
| 13 | 28 | M | White | Non-Hispanic | Case | 5 | 25 | 3 | 1.97 | 4.10 | 3.95 | 1.38 | 173.2 |
| 18 | 55 | M | White | Non-Hispanic | Case | 16 | 28 | 6 | 2.57 | 1.75 | 3.92 | 1.31 | 410.3 |
| 20 | 26 | F | White | Non-Hispanic | Case | 2 | 17 | 5 | 1.80 | 2.00 | 4.49 | 1.68 | 5.0 |
| 21 | 54 | M | White | Hispanic | Contact | 11 | 27 | 7 | 5.60 | 2.88 | 7.60 | 2.47 | 5052.7 |
| 24 | 34 | M | White | Non-Hispanic | Case | 15 | 30 | 4 | 2.29 | 1.97 | 4.36 | 1.45 | 280.7 |
| 26 | 66 | M | White | Non-Hispanic | Case | 2 | 35 | 3 | 1.67 | 1.86 | 4.00 | 1.58 | 276.1 |
| 27 | 28 | M | White | Non-Hispanic | Case | 9 | 32 | 4 | 1.79 | 1.64 | 4.41 | 1.30 | 739.3 |
| 28 | 26 | M | White | Non-Hispanic | Case | 7 | 21 | 4 | 2.39 | 2.25 | 4.93 | 1.66 | 888.9 |
| 29 | 26 | F | White | Non-Hispanic | Contact | 5 | 35 | 4 | 1.69 | 1.54 | 3.93 | 1.97 | 5.0 |
| 30 | 30 | M | White | Non-Hispanic | Contact | 3 | 35 | 4 | 3.07 | 2.55 | 4.86 | 1.34 | 5.0 |
| 31 | 51 | M | White | Non-Hispanic | Case | 9 | 33 | 3 | 1.29 | 1.60 | 4.69 | 1.20 | 192.3 |
| 32 | 46 | F | White | Non-Hispanic | Case | 8 | 32 | 3 | 1.30 | 2.18 | 3.79 | 0.83 | 47.4 |
| 37 | 27 | M | White | Non-Hispanic | Case | 6 | 28 | 6 | 2.13 | 1.52 | 4.11 | 1.04 | 286.3 |
| 38 | 57 | F | White | Non-Hispanic | Case | 10 | 38 | 4 | 1.87 | 2.02 | 4.30 | 1.12 | 518.9 |
| 40 | 44 | M | White | Non-Hispanic | Case | 7 | 23 | 5 | 1.72 | 1.50 | 3.81 | 1.47 | 42.1 |
| 41 | 35 | M | White | Non-Hispanic | Contact | 10 | 29 | 8 | 2.13 | 1.67 | 4.15 | 0.98 | 302.5 |
| 42 | 40 | M | White | Non-Hispanic | Contact | 20 | 36 | 8 | 2.37 | 1.50 | 5.82 | 1.43 | 627.1 |
| 45 | 55 | F | White | Non-Hispanic | Case | 7 | 48 | 6 | 1.95 | 0.97 | 3.53 | 0.75 | 5.0 |
| 46 | 39 | M | White | Non-Hispanic | Case | 8 | 30 | 2 | 2.68 | 1.45 | 4.14 | 1.29 | 59.2 |
| 47 | 43 | F | White | Non-Hispanic | Case | 11 | 33 | 5 | 3.02 | 2.30 | 6.23 | 2.66 | 10433.3 |
| 48 | 37 | F | White | Non-Hispanic | Case | 7 | 21 | 5 | 1.51 | 1.85 | 3.47 | 1.51 | 173.4 |
| 50 | 27 | F | White | Non-Hispanic | Contact | 7 | 28 | 4 | 1.64 | 2.32 | 5.42 | 1.45 | 924.7 |
| 51 | 21 | M | White | Non-Hispanic | Contact | 8 | 31 | 5 | 1.76 | 1.69 | 3.94 | 1.03 | 1499.2 |
| 54 | 40 | F | White | Non-Hispanic | Contact | 3 | 24 | 3 | 1.80 | 2.10 | 4.99 | 1.42 | 5.0 |
| 55 | 36 | M | White | Non-Hispanic | Case | 3 | 49 | 2 | 1.43 | 1.23 | 3.90 | 2.09 | 5.0 |
| 56 | 75 | F | White | Non-Hispanic | Case | 22 | 40 | 3 | 1.79 | 1.81 | 5.89 | 1.47 | 1388.4 |
| 57 | 66 | M | White | Non-Hispanic | Case | 6 | 21 | 5 | 1.54 | 2.10 | 4.33 | 1.00 | 2048.9 |
| 58 | 64 | F | White | Non-Hispanic | Contact | 1 | 32 | 2 | 1.20 | 1.71 | 3.95 | 1.21 | 5.0 |
| 64 | 28 | F | White | Non-Hispanic | Contact | 11 | 32 | 6 | 2.36 | 2.06 | 4.48 | 1.66 | 776.7 |
| 67 | 19 | F | N/A | Hispanic | Case | 5 | 29 | 6 | 2.56 | 1.98 | 5.48 | 1.32 | 2052.9 |
| 71 | 45 | F | White | Non-Hispanic | Case | 12 | 48 | 7 | 1.55 | 2.04 | 3.86 | 1.53 | 33.3 |
| 72 | 42 | M | White | Non-Hispanic | Case | 16 | 35 | 8 | 4.05 | 3.58 | 6.05 | 2.59 | 3138.2 |
| 75 | 46 | F | White | Non-Hispanic | Case | 10 | 36 | 4 | 1.64 | 2.37 | 3.98 | 1.20 | 271.5 |
| 76 | 49 | F | White | Non-Hispanic | Case | 28 | 34 | 4 | 1.88 | 1.65 | 5.17 | 0.97 | 219.8 |
| 77 | 37 | M | White | Non-Hispanic | Contact | 6 | 33 | 4 | 1.33 | 1.83 | 3.56 | 1.16 | 5.0 |
| 81 | 44 | F | White | Non-Hispanic | Contact | 3 | 35 | 2 | 1.58 | 1.73 | 3.82 | 1.10 | 5.0 |
| 82 | 46 | M | N/A | Non-Hispanic | Case | 0 |  | 0 | 2.19 | 1.92 | 5.41 | 1.81 | 130.7 |
| 88 | 41 | M | White | Non-Hispanic | Case | 7 | 23 | 4 | 1.82 | 3.32 | 4.97 | 1.37 | 424.7 |
| 95 | 44 | M | White | Non-Hispanic | Case | 9 | 36 | 6 | 2.61 | 1.62 | 6.03 | 1.62 | 961.9 |
| 96 | 48 | F | White | Non-Hispanic | Case | 9 | 30 | 3 | 3.93 | 1.93 | 6.25 | 2.26 | 927.7 |
| 97 | 39 | M | White | Non-Hispanic | Case | 9 | 31 | 3 | 1.58 | 1.61 | 4.03 | 2.33 | 202.7 |
| 98 | 35 | F | White | Non-Hispanic | Case | 2 | 24 | 4 | 1.78 | 1.47 | 5.38 | 1.22 | 249.0 |
| 99 | 36 | F | White | Non-Hispanic | Case | 13 | 29 | 5 | 2.50 | 3.27 | 4.38 | 2.49 | 1127.6 |
| 107 | 53 | F | White | Non-Hispanic | Contact | 10 | 29 | 4 | 1.74 | 1.41 | 4.66 | 0.90 | 297.5 |
| 108 | 75 | M | White | Non-Hispanic | Case | 16 | 41 | 7 | 1.37 | 0.88 | 3.47 | 1.36 | 557.5 |
| 110 | 27 | M | White | Non-Hispanic | Case | 1 | 25 | 1 | 1.30 | 1.60 | 4.00 | 0.93 | 5.0 |
| 114 | 30 | F | White | Non-Hispanic | Case | 15 | 36 | 7 | 1.65 | 1.92 | 3.53 | 1.55 | 110.9 |
| 115 | 65 | F | White | Non-Hispanic | Contact | 20 | 41 | 6 | 2.10 | 3.28 | 4.32 | 3.27 | 1127.7 |
| 119 | 56 | M | White | Non-Hispanic | Case | 13 | 48 | 3 | 1.36 | 1.26 | 4.41 | 1.59 | 650.3 |
| 120 | 56 | F | White | Non-Hispanic | Case | 26 | 48 | 6 | 0.99 | 0.97 | 3.65 | 1.21 | 100.6 |
| 121 | 19 | M | White | Non-Hispanic | Contact | 3 | 42 | 2 | 0.85 | 0.87 | 3.56 | 1.26 | 5.0 |
| 122 | 21 | F | White | Non-Hispanic | Contact | 3 | 38 | 1 | 1.44 | 0.94 | 3.39 | 1.22 | 5.0 |
| 123 | 26 | M | White | Non-Hispanic | Contact | 12 | 34 | 6 | 0.94 | 0.95 | 3.39 | 1.40 | 5.0 |
| 124 | 63 | F | Asian | Non-Hispanic | Contact | 4 | 37 | 3 | 1.58 | 2.06 | 3.49 | 1.32 | 5.0 |
| 125 | 51 | F | White | Non-Hispanic | Case | 10 | 26 | 3 | 1.92 | 3.49 | 3.86 | 1.24 | 126.5 |
| 127 | 24 | F | White | Non-Hispanic | Case | 10 | 43 | 6 | 1.80 | 2.50 | 4.37 | 2.41 | 883.5 |
| 130 | 39 | M | White | Non-Hispanic | Contact | 7 | 28 | 5 | 1.24 | 1.72 | 3.91 | 1.34 | 5.0 |
| 131 | 39 | M | White | Non-Hispanic | Case | 5 | 25 | 4 | 1.46 | 1.38 | 4.44 | 1.03 | 7.8 |
| 132 | 36 | M | White | Non-Hispanic | Contact | 10 | 50 | 6 | 2.15 | 1.76 | 4.84 | 1.97 | 5.0 |
| 134 | 27 | F | White | Non-Hispanic | Contact | 16 | 22 | 5 | 2.51 | 2.18 | 6.84 | 1.94 | 2700.6 |
| 135 | 62 | F | White | Non-Hispanic | Case | 8 | 31 | 6 | 2.20 | 2.02 | 3.80 | 1.13 | 350.0 |
| 140 | 63 | F | White | Non-Hispanic | Case | 28 | 47 | 1 | 1.05 | 1.24 | 3.58 | 1.28 | 52.4 |
| 149 | 41 | M | White | Non-Hispanic | Contact | 17 | 28 | 6 | 1.68 | 2.02 | 3.67 | 1.09 | 494.9 |
| 150 | 50 | F | White | Non-Hispanic | Contact | 12 | 45 | 7 | 1.15 | 0.43 | 3.18 | 1.82 | 5.0 |
| 154 | 68 | M | Asian | Non-Hispanic | Case | 16 | 30 | 9 | 3.19 | 2.19 | 4.85 | 1.29 | 928.2 |
| 157 | 50 | M | White | Non-Hispanic | Case | 10 | 32 | 8 | 2.40 | 2.86 | 3.90 | 2.06 | 741.7 |
| 166 | 28 | F | White | Non-Hispanic | Case | 13 | 45 | 2 | 1.27 | 0.66 | 3.45 | 0.94 | 5.0 |
| 167 | 50 | F | White | Non-Hispanic | Contact | 11 | 41 | 6 | 1.43 | 3.71 | 3.93 | 8.74 | 5.0 |
| 172 | 38 | F | White | Non-Hispanic | Case | 8 | 22 | 9 | 1.71 | 2.58 | 4.29 | 1.20 | 301.1 |
| 173 | 47 | M | White | Non-Hispanic | Case | 5 | 47 | 7 | 2.57 | 4.14 | 4.78 | 4.48 | 646.9 |
| 178 | 26 | F | White | Non-Hispanic | Case | 6 | 24 | 4 | 1.54 | 1.59 | 3.66 | 1.02 | 5.0 |
| 179 | 39 | M | White | Non-Hispanic | Contact | 10 | 37 | 3 | 1.89 | 2.25 | 3.83 | 1.73 | 370.1 |
| 182 | 44 | F | White | Non-Hispanic | Contact | 10 | 38 | 6 | 3.80 | 1.77 | 5.36 | 8.05 | 1503.7 |
| 183 | 43 | F | White | Non-Hispanic | Case | 13 | 44 | 8 | 1.56 | 1.02 | 3.88 | 1.55 | 240.1 |
| 185 | 54 | M | White | Non-Hispanic | Case | 11 | 44 | 8 | 3.51 | 1.39 | 5.55 | 2.11 | 1806.8 |
| 186 | 38 | F | N/A | N/A | Case | 8 | 26 | 2 | 1.73 | 2.47 | 4.37 | 1.23 | 296.9 |
| 190 | 54 | F | White | Non-Hispanic | Case | 18* | 63 | 9 | 3.24 | 1.24 | 7.38 | 1.42 | 598.1 |
| 195 | 24 | M | White | Non-Hispanic | Case | 18 | 42 | 5 | 2.74 | 2.55 | 5.01 | 2.33 | 1315.1 |
| 200 | 60 | F | White | Non-Hispanic | Case | 17 | 39 | 7 | 2.40 | 1.00 | 4.38 | 1.51 | 1014.4 |
| 201 | 50 | M | White | Non-Hispanic | Contact | 15 | 33 | 6 | 4.37 | 2.57 | 6.15 | 1.57 | 3897.4 |
| 202 | 57 | M | White | Non-Hispanic | Case | 21 | 34 | 7 | 2.10 | 2.08 | 5.07 | 1.17 | 257.9 |
| 205 | 64 | M | White | Non-Hispanic | Case | 7 | 36 | 4 | 4.51 | 0.70 | 6.12 | 1.69 | 924.4 |
| 222 | 28 | M | Asian | Non-Hispanic | Case | 11 | 29 | 7 | 1.28 | 0.69 | 3.94 | 3.46 | 5.0 |
| 229 | 45 | M | White | Non-Hispanic | Case | 10 | 63 | 4 | 2.92 | 1.42 | 4.90 | 1.58 | 1272.9 |
| 230 | 50 | M | White | Non-Hispanic | Case | 18 | 33 | 7 | 3.80 | 0.47 | 3.48 | 0.88 | 5.0 |
| 232 | 38 | F | White | Non-Hispanic | Case | 13 | 43 | 7 | 1.57 | 0.70 | 4.24 | 5.70 | 94.3 |
| 233 | 55 | M | White | Non-Hispanic | Case | 20 | 41 | 3 | 2.07 | 2.11 | 4.51 | 1.07 | 173.2 |
| 241 | 36 | M | White | Non-Hispanic | Case | 12 | 30 | 7 | 2.27 | 2.66 | 4.54 | 1.46 | 923.1 |
| 242 | 59 | M | White | Non-Hispanic | Case | 10 | 42 | 6 | 4.91 | 1.94 | 4.81 | 2.16 | 1353.0 |
| 243 | 30 | F | Asian | Non-Hispanic | Case | 6 | 26 | 5 | 2.92 | 2.57 | 5.06 | 1.14 | 1300.2 |
| 246 | 44 | F | White | Non-Hispanic | Case | 10 | 38 | 7 | 2.05 | 2.79 | 6.09 | 1.32 | 566.0 |
| 255 | 33 | M | White | Non-Hispanic | Case | 14 | 44 | 6 | 2.14 | 0.70 | 4.20 | 1.24 | 172.5 |
| 256 | 63 | F | White | Non-Hispanic | Case | 27 | 42 | 6 | 1.72 | 1.96 | 4.26 | 7.79 | 141.6 |
| 258 | 52 | M | White | Non-Hispanic | Contact | 14 | 48 | 6 | 2.64 | 1.20 | 4.52 | 1.85 | 4145.9 |
| 279 | 41 | M | White | Non-Hispanic | Case | 7 | 38 | 8 | 1.68 | 2.13 | 3.77 | 1.90 | 308.9 |
| 280 | 59 | M | White | Non-Hispanic | Case | 6 | 32 | 7 | 2.53 | 3.07 | 4.61 | 1.19 | 1072.1 |
| 302 | 47 | F | White | Non-Hispanic | Case | 35* | 49 | 7 | 1.48 | 0.97 | 4.06 | 1.26 | 5.0 |
| 310 | 34 | F | White | Non-Hispanic | Case | 17 | 35 | 5 | 3.95 | 1.24 | 9.44 | 3.07 | 485.5 |
| 314 | 46 | M | White | Non-Hispanic | Case | 11 | 38 | 7 | 2.12 | 0.88 | 4.56 | 1.51 | 667.1 |
| 315 | 29 | F | White | Non-Hispanic | Case | 15 | 42 | 8 | 3.02 | 0.69 | 3.98 | 1.03 | 376.5 |
| 319 | 50 | M | White | Non-Hispanic | Case | 5 | 38 | 6 | 3.71 | 2.28 | 3.79 | 1.05 | 5.0 |
| 323 | 39 | F | White | Non-Hispanic | Case | 7 | 45 | 7 | 1.05 | 1.03 | 3.53 | 1.42 | 5.0 |
| 325 | 52 | M | White | Non-Hispanic | Case | 16 | 38 | 8 | 2.25 | 1.47 | 4.83 | 2.28 | 1603.3 |
| 343 | 21 | F | White | Non-Hispanic | Case | 16 | 49 | 5 | 1.63 | 0.94 | 3.37 | 1.70 | 5.0 |
| 352 | 44 | M | White | Non-Hispanic | Case | 16 | 43 | 4 | 3.54 | 0.92 | 4.50 | 1.07 | 519.2 |
| 353 | 60 | M | White | Non-Hispanic | Case | 14 | 49 | 6 | 5.38 | 1.12 | 5.69 | 1.05 | 855.5 |
| 356 | 22 | F | White | Non-Hispanic | Contact | 16 | 38 | 3 | 1.37 | 0.61 | 2.98 | 1.09 | 5.0 |
| 357 | 27 | F | White | Non-Hispanic | Contact | 34 | 56 | 5 | 7.49 | 1.10 | 2.77 | 1.13 | 5.0 |
| 364 | 29 | M | White | Non-Hispanic | Contact | 14 | 49 | 6 | 0.97 | 0.58 | 2.90 | 0.89 | 5.0 |
| 366 | 41 | F | White | Non-Hispanic | Contact | 9 | 34 | 7 | 0.98 | 0.52 | 3.51 | 1.19 | 5.0 |
| 373 | 35 | F | White | Non-Hispanic | Case | 12 | 51 | 7 | 1.69 | 1.26 | 5.14 | 1.69 | 5.0 |
| 388 | 47 | F | White | Non-Hispanic | Contact | 14 | 41 | 9 | 1.57 | 1.06 | 3.61 | 1.69 | 5.0 |
| 393 | 69 | M | White | Non-Hispanic | Case | 23* | 54 | 9 | 1.28 | 1.74 | 3.81 | 2.65 | 715.4 |
| 394 | 48 | F | Multiple | Hispanic | Case | 7 | 67 | 4 | 2.05 | 0.87 | 4.34 | 2.02 | 1281.5 |
| 397 | 52 | M | White | Non-Hispanic | Case | 22 | 45 | 8 | 3.32 | 0.59 | 5.01 | 0.87 | 1516.9 |
| 403 | 52 | M | Asian | Non-Hispanic | Case | 18* | 39 | 10 | 5.36 | 1.09 | 10.01 | 1.36 | 3887.8 |
| 406 | 65 | M | White | Non-Hispanic | Case | 20 | 56 | 8 | 4.69 | 0.90 | 7.51 | 1.15 | 1288.7 |
| 410 | 34 | M | White | Non-Hispanic | Case | 12 | 46 | 8 | 1.06 | 0.56 | 3.95 | 0.76 | 5.0 |
| 421 | 62 | F | White | Non-Hispanic | Contact | 12 | 43 | 9 | 0.95 | 1.07 | 3.34 | 1.35 | 5.0 |
| 426 | 65 | M | White | Non-Hispanic | Case | 18 | 51 | 6 | 2.07 | 0.55 | 3.95 | 1.49 | 804.8 |
| 437 | 43 | F | Asian | Non-Hispanic | Case | 14 | 34 | 7 | 2.54 | 0.47 | 4.30 | 1.44 | 698.8 |
| 460 | 36 | M | White | Non-Hispanic | Case | 11 | 39 | 6 | 2.94 | 3.18 | 5.51 | 2.80 | 1906.7 |
| 461 | 49 | M | White | Non-Hispanic | Case | 7 | 39 | 5 | 3.38 | 0.94 | 4.67 | 2.02 | 1076.6 |
| 462 | 28 | F | White | Non-Hispanic | Case | 16 | 45 | 5 | 1.36 | 0.38 | 3.07 | 1.11 | 5.0 |
| 470 | 28 | F | White | Non-Hispanic | Case | 17 | 51 | 4 | 1.26 | 0.86 | 3.97 | 1.50 | 5.0 |
| 478 | 31 | M | White | Non-Hispanic | Case | 16 | 52 | 4 | 1.43 | 0.93 | 3.70 | 1.97 | 263.2 |
| 481 | 28 | F | Asian | Non-Hispanic | Case | 15 | 43 | 8 | 1.70 | 0.39 | 3.46 | 1.24 | 5.0 |
| 486 | 64 | F | White | Non-Hispanic | Case | 11 | 41 | 10 | 1.70 | 1.00 | 3.68 | 1.29 | 5.0 |
| 500 | 46 | M | White | Non-Hispanic | Case | 12 | 53 | 5 | 1.10 | 0.82 | 3.49 | 1.34 | 5.0 |
| 501 | 32 | M | Asian | Non-Hispanic | Case | 18* | 53 | 10 | 2.62 | 0.65 | 4.51 | 1.21 | 718.8 |
| 502 | 52 | M | White | Non-Hispanic | Case | 16* | 53 | 9 | 5.10 | 0.61 | 5.10 | 1.62 | 2171.8 |
| 506 | 46 | M | White | Non-Hispanic | Case | 12 | 59 | 9 | 0.84 | 0.81 | 3.13 | 1.21 | 5.0 |
| 507 | 39 | M | White | Non-Hispanic | Case | 15 | 60 | 8 | 1.92 | 0.96 | 4.56 | 1.52 | 5.0 |
| 509 | 36 | M | White | Non-Hispanic | Case | 11 | 50 | 5 | 1.99 | 1.01 | 3.99 | 1.45 | 5.0 |
| 526 | 49 | M | Asian | Non-Hispanic | Case | 11 | 34 | 7 | 3.36 | 1.45 | 5.88 | 1.57 | 4193.3 |
| 537 | 52 | M | White | Non-Hispanic | Case | 15 | 45 | 6 | 1.47 | 0.95 | 3.65 | 1.58 | 923.3 |
| 539 | 73 | F | White | Non-Hispanic | Case | 19* | 54 | 10 | 2.82 | 0.63 | 4.46 | 1.45 | 487.9 |
| 547 | 59 | M | White | Non-Hispanic | Case | 15* | 36 | 9 | 2.97 | 1.53 | 5.08 | 2.59 | 2900.6 |
| 587 | 54 | M | PI | N/A | Case | 17* | 51 | 8 | 3.22 | 0.60 | 4.01 | 1.49 | 473.1 |
| 632 | 38 | M | White | Non-Hispanic | Contact | 10 | 43 | 6 | 2.49 | 0.86 | 4.50 | 1.63 | 572.3 |
| 633 | 39 | M | White | Non-Hispanic | Contact | 8 | 57 | 4 | 1.25 | 1.04 | 3.38 | 1.73 | 5.0 |
| 652 | 76 | M | White | Non-Hispanic | Case | 18* | 56 | 10 | 4.75 | 1.46 | 8.96 | 3.80 | 2324.0 |
| 664 | 45 | F | White | Non-Hispanic | Case | 17* | 42 | 10 | 1.68 | 0.43 | 3.93 | 1.32 | 5.0 |
| 675 | 47 | M | White | Non-Hispanic | Contact | 31 | 47 | 5 | 0.79 | 0.93 | 2.94 | 1.38 | 5.0 |

\*hospitalized, Sx=symptoms

**Extended Data Table 2.**
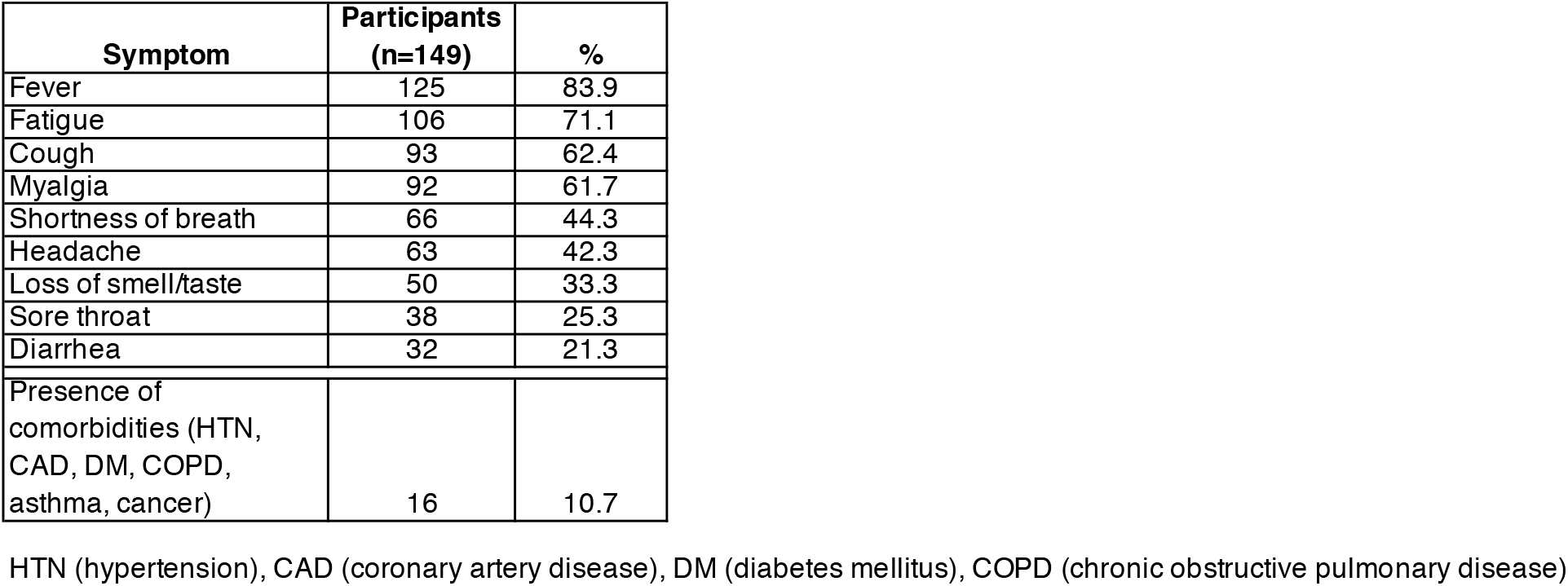
Frequency of symptoms and comorbidities reported by participants

| **Symptom** | **Participants**<br>**(n=149)** | **%** |
| --- | --- | --- |
| Fever | 125 | 83.9 |
| Fatigue | 106 | 71.1 |
| Cough | 93 | 62.4 |
| Myalgia | 92 | 61.7 |
| Shortness of breath | 66 | 44.3 |
| Headache | 63 | 42.3 |
| Loss of smell/taste | 50 | 33.3 |
| Sore throat | 38 | 25.3 |
| Diarrhea | 32 | 21.3 |
| Presence of<br>comorbidities (HTN,<br>CAD, DM, COPD,<br>asthma, cancer) | 16 | 10.7 |

HTN (hypertension), CAD (coronary artery disease), DM (diabetes mellitus), COPD (chronic obstructive pulmonary disease)

**Extended Data Tables 3 and 4 are provided as separate Excel files.**

**Extended Data Table 5.**
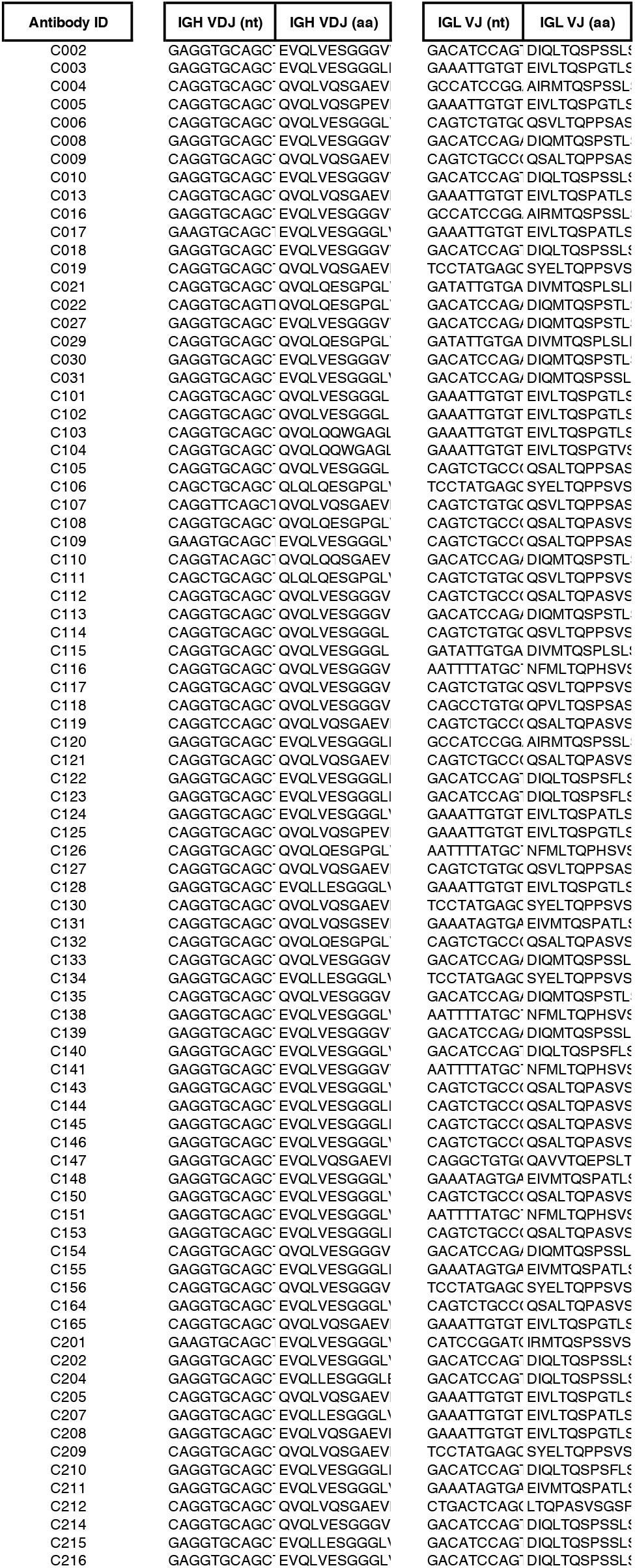
Sequences of cloned recombinant antibodies

**Extended Data Table 6.**
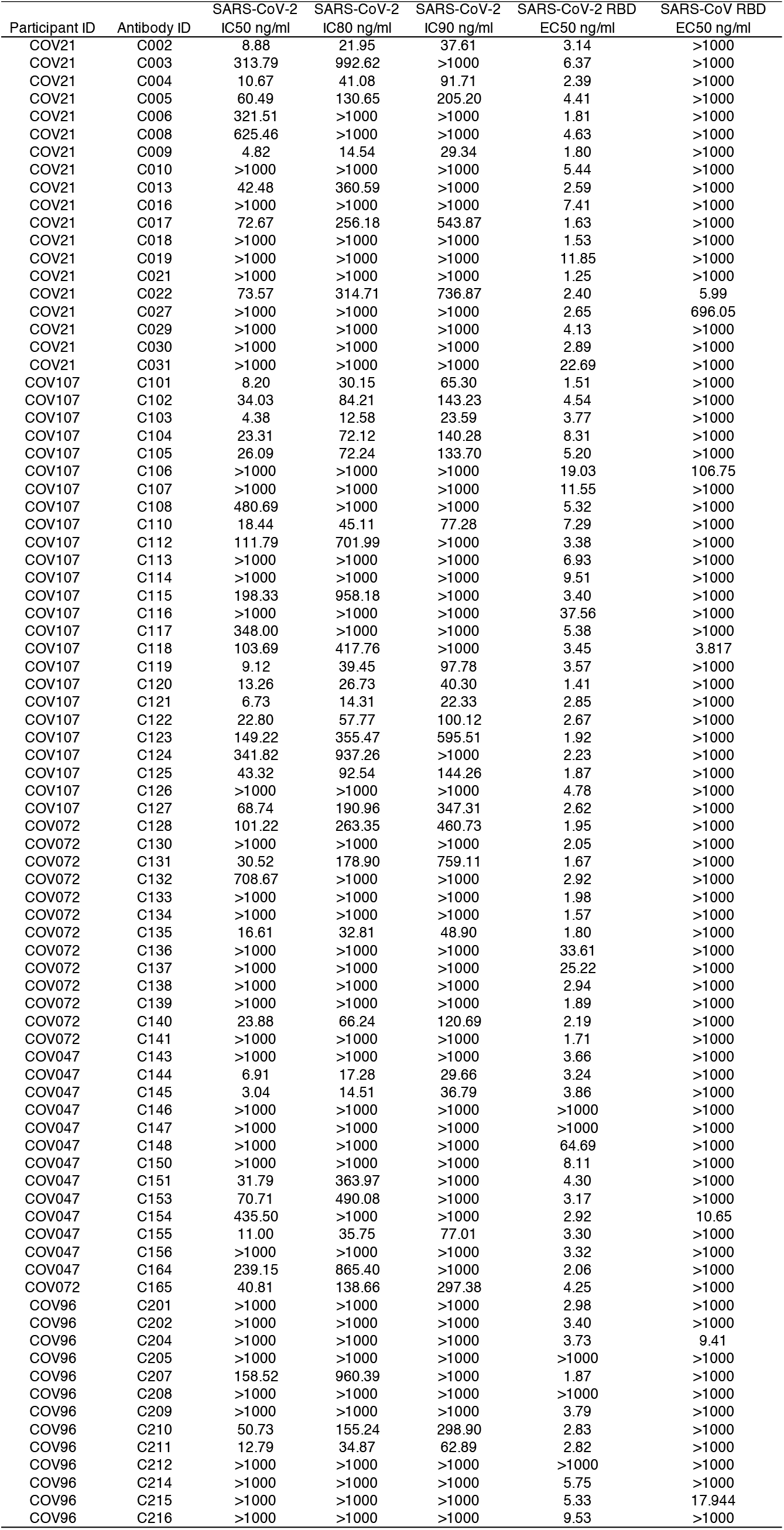
Effective and inhibitory concentrations of the monoclonal antibodies

| Participant ID | Antibody ID | SARS-CoV-2 | SARS-CoV-2 | SARS-CoV-2 | SARS-CoV-2 RBD | SARS-CoV RBD |
| --- | --- | --- | --- | --- | --- | --- |
|  |  | IC50 ng/ml | IC80 ng/ml | IC90 ng/ml | EC50 ng/ml | EC50 ng/ml |
| COV21 | C002 | 8.88 | 21.95 | 37.61 | 3.14 | >1000 |
| COV21 | C003 | 313.79 | 992.62 | >1000 | 6.37 | >1000 |
| COV21 | C004 | 10.67 | 41.08 | 91.71 | 2.39 | >1000 |
| COV21 | C005 | 60.49 | 130.65 | 205.20 | 4.41 | >1000 |
| COV21 | C006 | 321.51 | >1000 | >1000 | 1.81 | >1000 |
| COV21 | C008 | 625.46 | >1000 | >1000 | 4.63 | >1000 |
| COV21 | C009 | 4.82 | 14.54 | 29.34 | 1.80 | >1000 |
| COV21 | C010 | >1000 | >1000 | >1000 | 5.44 | >1000 |
| COV21 | C013 | 42.48 | 360.59 | >1000 | 2.59 | >1000 |
| COV21 | C016 | >1000 | >1000 | >1000 | 7.41 | >1000 |
| COV21 | C017 | 72.67 | 256.18 | 543.87 | 1.63 | >1000 |
| COV21 | C018 | >1000 | >1000 | >1000 | 1.53 | >1000 |
| COV21 | C019 | >1000 | >1000 | >1000 | 11.85 | >1000 |
| COV21 | C021 | >1000 | >1000 | >1000 | 1.25 | >1000 |
| COV21 | C022 | 73.57 | 314.71 | 736.87 | 2.40 | 5.99 |
| COV21 | C027 | >1000 | >1000 | >1000 | 2.65 | 696.05 |
| COV21 | C029 | >1000 | >1000 | >1000 | 4.13 | >1000 |
| COV21 | C030 | >1000 | >1000 | >1000 | 2.89 | >1000 |
| COV21 | C031 | >1000 | >1000 | >1000 | 22.69 | >1000 |
| COV107 | C101 | 8.20 | 30.15 | 65.30 | 1.51 | >1000 |
| COV107 | C102 | 34.03 | 84.21 | 143.23 | 4.54 | >1000 |
| COV107 | C103 | 4.38 | 12.58 | 23.59 | 3.77 | >1000 |
| COV107 | C104 | 23.31 | 72.12 | 140.28 | 8.31 | >1000 |
| COV107 | C105 | 26.09 | 72.24 | 133.70 | 5.20 | >1000 |
| COV107 | C106 | >1000 | >1000 | >1000 | 19.03 | 106.75 |
| COV107 | C107 | >1000 | >1000 | >1000 | 11.55 | >1000 |
| COV107 | C108 | 480.69 | >1000 | >1000 | 5.32 | >1000 |
| COV107 | C110 | 18.44 | 45.11 | 77.28 | 7.29 | >1000 |
| COV107 | C112 | 111.79 | 701.99 | >1000 | 3.38 | >1000 |
| COV107 | C113 | >1000 | >1000 | >1000 | 6.93 | >1000 |
| COV107 | C114 | >1000 | >1000 | >1000 | 9.51 | >1000 |
| COV107 | C115 | 198.33 | 958.18 | >1000 | 3.40 | >1000 |
| COV107 | C116 | >1000 | >1000 | >1000 | 37.56 | >1000 |
| COV107 | C117 | 348.00 | >1000 | >1000 | 5.38 | >1000 |
| COV107 | C118 | 103.69 | 417.76 | >1000 | 3.45 | 3.817 |
| COV107 | C119 | 9.12 | 39.45 | 97.78 | 3.57 | >1000 |
| COV107 | C120 | 13.26 | 26.73 | 40.30 | 1.41 | >1000 |
| COV107 | C121 | 6.73 | 14.31 | 22.33 | 2.85 | >1000 |
| COV107 | C122 | 22.80 | 57.77 | 100.12 | 2.67 | >1000 |
| COV107 | C123 | 149.22 | 355.47 | 595.51 | 1.92 | >1000 |
| COV107 | C124 | 341.82 | 937.26 | >1000 | 2.23 | >1000 |
| COV107 | C125 | 43.32 | 92.54 | 144.26 | 1.87 | >1000 |
| COV107 | C126 | >1000 | >1000 | >1000 | 4.78 | >1000 |
| COV107 | C127 | 68.74 | 190.96 | 347.31 | 2.62 | >1000 |
| COV072 | C128 | 101.22 | 263.35 | 460.73 | 1.95 | >1000 |
| COV072 | C130 | >1000 | >1000 | >1000 | 2.05 | >1000 |
| COV072 | C131 | 30.52 | 178.90 | 759.11 | 1.67 | >1000 |
| COV072 | C132 | 708.67 | >1000 | >1000 | 2.92 | >1000 |
| COV072 | C133 | >1000 | >1000 | >1000 | 1.98 | >1000 |
| COV072 | C134 | >1000 | >1000 | >1000 | 1.57 | >1000 |
| COV072 | C135 | 16.61 | 32.81 | 48.90 | 1.80 | >1000 |
| COV072 | C136 | >1000 | >1000 | >1000 | 33.61 | >1000 |
| COV072 | C137 | >1000 | >1000 | >1000 | 25.22 | >1000 |
| COV072 | C138 | >1000 | >1000 | >1000 | 2.94 | >1000 |
| COV072 | C139 | >1000 | >1000 | >1000 | 1.89 | >1000 |
| COV072 | C140 | 23.88 | 66.24 | 120.69 | 2.19 | >1000 |
| COV072 | C141 | >1000 | >1000 | >1000 | 1.71 | >1000 |
| COV047 | C143 | >1000 | >1000 | >1000 | 3.66 | >1000 |
| COV047 | C144 | 6.91 | 17.28 | 29.66 | 3.24 | >1000 |
| COV047 | C145 | 3.04 | 14.51 | 36.79 | 3.86 | >1000 |
| COV047 | C146 | >1000 | >1000 | >1000 | >1000 | >1000 |
| COV047 | C147 | >1000 | >1000 | >1000 | >1000 | >1000 |
| COV047 | C148 | >1000 | >1000 | >1000 | 64.69 | >1000 |
| COV047 | C150 | >1000 | >1000 | >1000 | 8.11 | >1000 |
| COV047 | C151 | 31.79 | 363.97 | >1000 | 4.30 | >1000 |
| COV047 | C153 | 70.71 | 490.08 | >1000 | 3.17 | >1000 |
| COV047 | C154 | 435.50 | >1000 | >1000 | 2.92 | 10.65 |
| COV047 | C155 | 11.00 | 35.75 | 77.01 | 3.30 | >1000 |
| COV047 | C156 | >1000 | >1000 | >1000 | 3.32 | >1000 |
| COV047 | C164 | 239.15 | 865.40 | >1000 | 2.06 | >1000 |
| COV072 | C165 | 40.81 | 138.66 | 297.38 | 4.25 | >1000 |
| COV96 | C201 | >1000 | >1000 | >1000 | 2.98 | >1000 |
| COV96 | C202 | >1000 | >1000 | >1000 | 3.40 | >1000 |
| COV96 | C204 | >1000 | >1000 | >1000 | 3.73 | 9.41 |
| COV96 | C205 | >1000 | >1000 | >1000 | >1000 | >1000 |
| COV96 | C207 | 158.52 | 960.39 | >1000 | 1.87 | >1000 |
| COV96 | C208 | >1000 | >1000 | >1000 | >1000 | >1000 |
| COV96 | C209 | >1000 | >1000 | >1000 | 3.79 | >1000 |
| COV96 | C210 | 50.73 | 155.24 | 298.90 | 2.83 | >1000 |
| COV96 | C211 | 12.79 | 34.87 | 62.89 | 2.82 | >1000 |
| COV96 | C212 | >1000 | >1000 | >1000 | >1000 | >1000 |
| COV96 | C214 | >1000 | >1000 | >1000 | 5.75 | >1000 |
| COV96 | C215 | >1000 | >1000 | >1000 | 5.33 | 17.944 |
| COV96 | C216 | >1000 | >1000 | >1000 | 9.53 | >1000 |

## Extended Data Figures

**Extended Data Figure 1.**
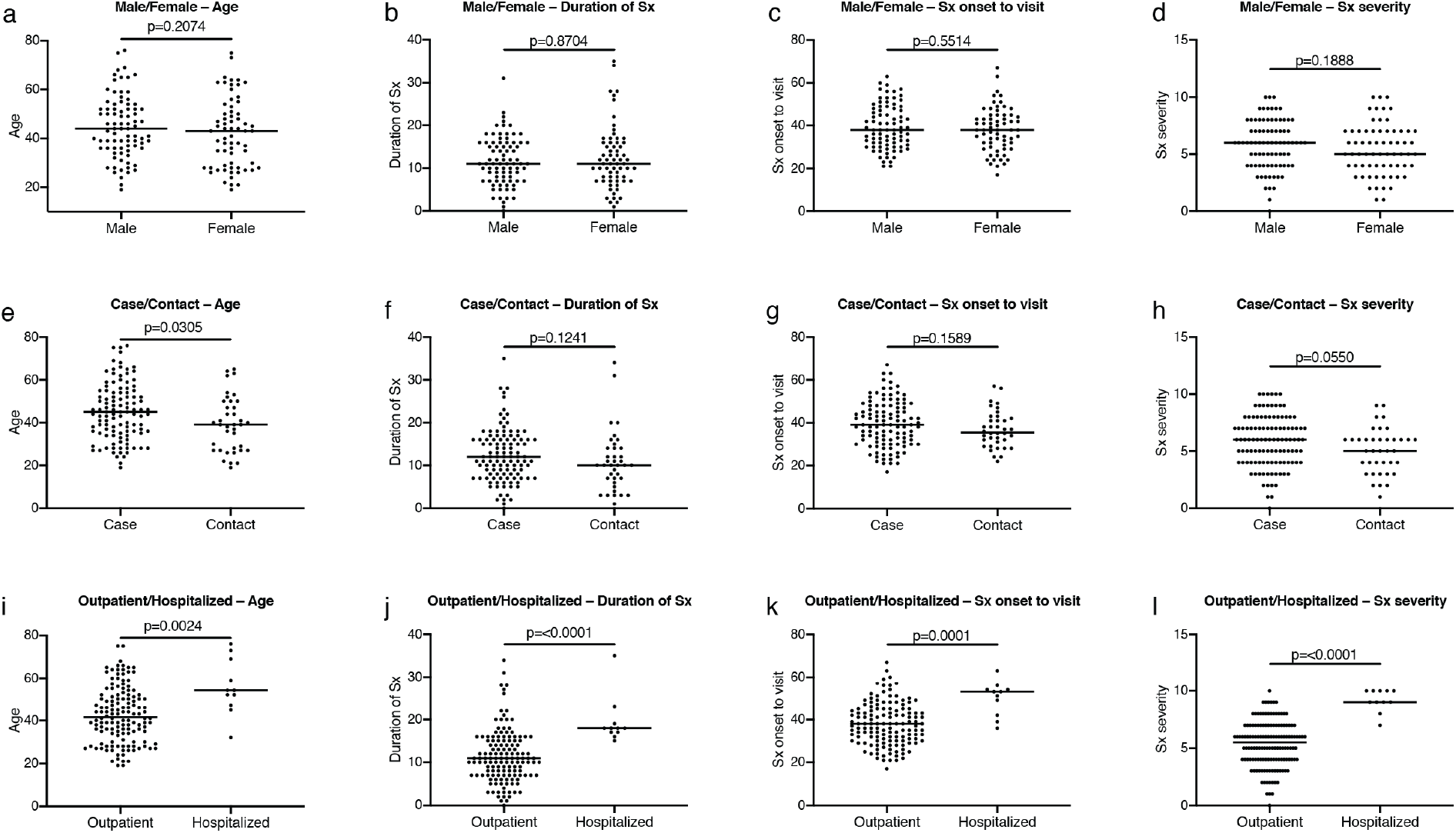
Clinical correlates. **a,** Age distribution (Y axis) for all males and females in the cohort p=0.2074. **b,** Duration of symptoms in days (Y axis) for all males and females in the cohort p=0.8704. **c,** Time between symptom onset and plasma collection (Y axis) for all males and females in the cohort p=0.5514. **d,** Subjective symptom severity on a scale of 0-10 (Y axis) for all males and females in the cohort p=0.1888. **e,** Age distribution (Y axis) for all cases and contacts in the cohort p=0.0305. **f,** Duration of symptoms in days (Y axis) for all cases and contacts in the cohort p=0.1241. **g,** Time between symptom onset and plasma collection in days (Y axis) for all cases and contacts in the cohort p=0.1589. **h,** Symptom severity (Y axis) for all cases and contacts in the cohort p=0.0550. **i,** Age distribution (Y axis) for all outpatient and hospitalized participants p=0.0024. **j,** Duration of symptoms in days (Y axis) for all outpatient and hospitalized participants in the cohort p=<0.0001. **k,** Time between symptom onset and plasma collection in days (Y axis) for all outpatient and hospitalized participants in the cohort p=0.0001. **l,** Symptom severity (Y axis) for all outpatient and hospitalized participants in the cohort p=<0.0001. Horizontal bars indicate median values. Statistical significance was determined using two-tailed Mann-Whitney U test.

**Extended Data Figure 2.**
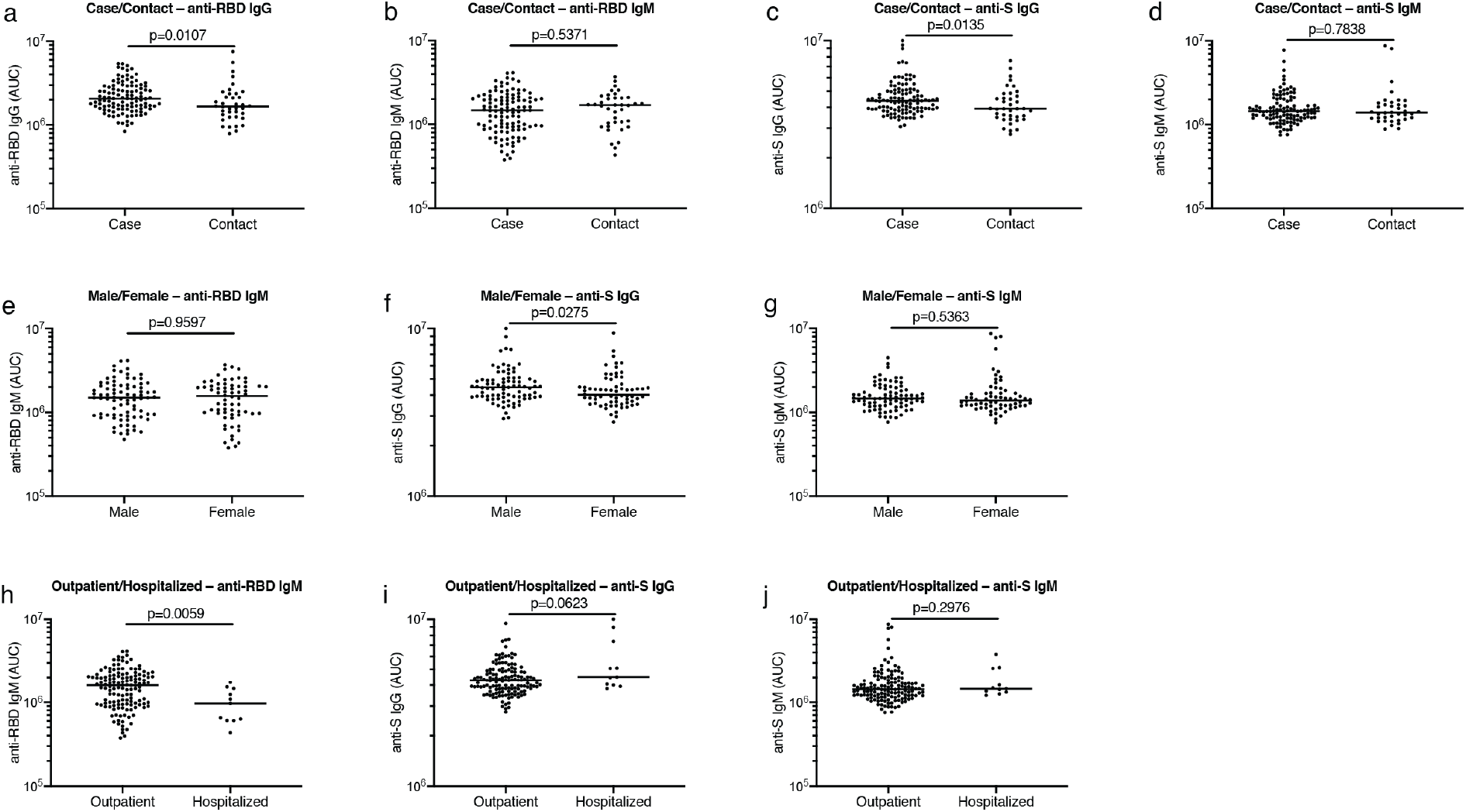
Clinical correlates of plasma antibody titers. **a,** AUC for IgG anti-RBD (Y axis) for all cases and contacts in the cohort p=0.0107. **b,** AUC for IgM anti-RBD (Y axis) for all cases and contacts in the cohort p=0.5371. **c,** AUC for IgG anti-S (Y axis) for all cases and contacts in the cohort p=0.0135. **d,** AUC for IgM anti-S (Y axis) for all cases and contacts in the cohort p=0.7838. **e,** AUC for IgM anti-RBD (Y axis) for all males and females in the cohort p=0.9597. **f,** AUC for IgG anti-S (Y axis) for all males and females in the cohort p=0.0275. **g,** AUC for IgM anti-S (Y axis) for all males and females in the cohort p=0.5363. **h,** AUC for IgM anti-RBD (Y axis) for all outpatient and hospitalized participants in the cohort p=0.0059. **i,** AUC for IgG anti-S (Y axis) for all outpatient and hospitalized participants in the cohort p=0.0623. **j,** AUC for IgM anti-S (Y axis) for all outpatient and hospitalized participants in the cohort p=0.2976. Horizontal bars indicate median values. Statistical significance was determined using two-tailed Mann-Whitney U test.

**Extended Data Figure 3.**
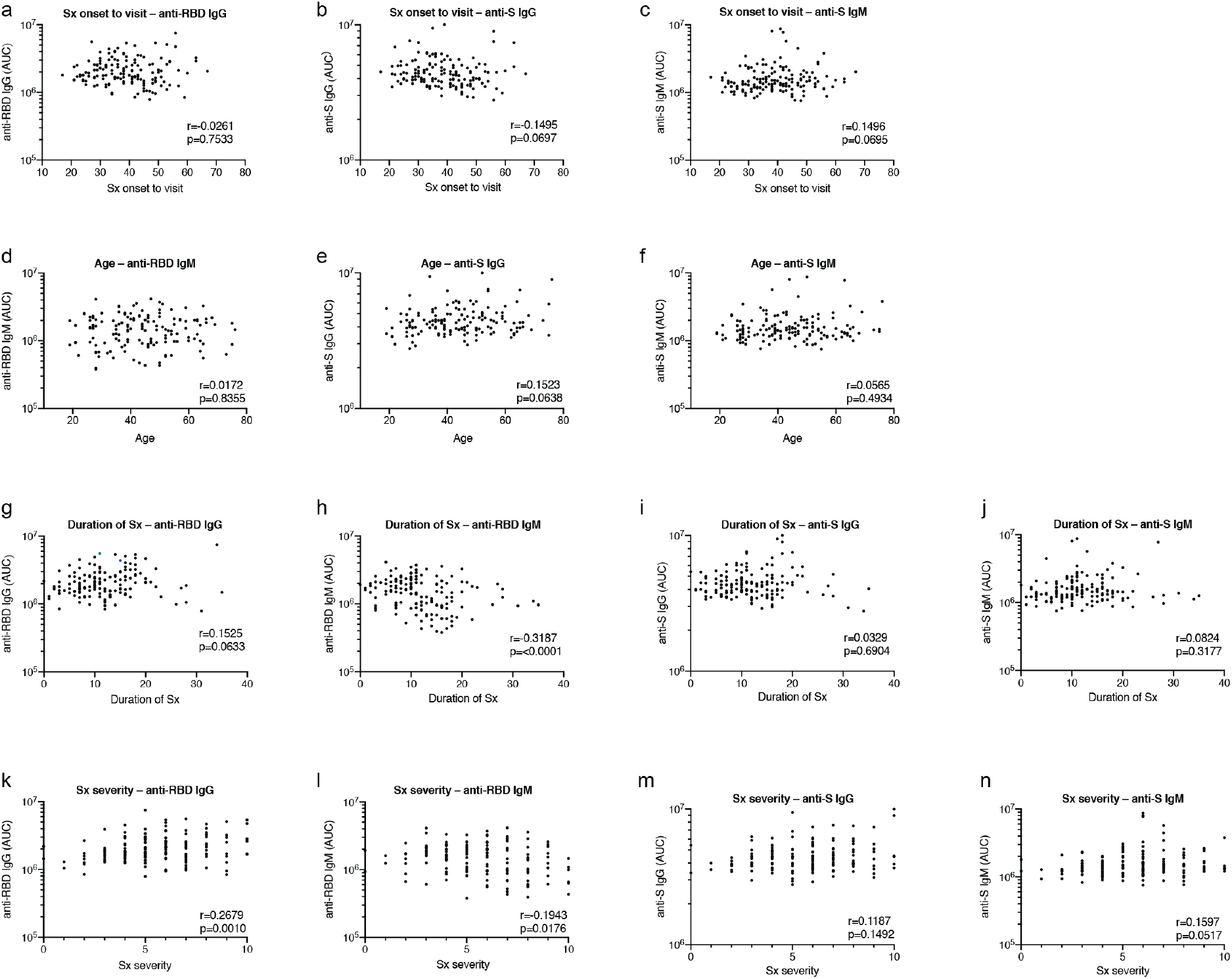
Additional clinical correlates of plasma antibody titers. **a**, Time between symptom onset and plasma collection in days (X axis) plotted against AUC for IgG anti-RBD (Y axis) r=-0.0261 p=0.7533. **b**, Time between symptom onset and plasma collection in days (X axis) plotted against AUC for IgG anti-S (Y axis) r=-0.1495 p=0.0697. **c**, Time between symptom onset and plasma collection in days (X axis) plotted against AUC for IgM anti-S (Y axis) r=0.1496 p=0.0695. **d**, Age (X axis) plotted against AUC for IgM anti-RBD (Y axis) r=0.0172 p=0.8355. **e**, Age (X axis) plotted against AUC for IgG anti-S (Y axis) r=0.1523 p=0.0638. **f**, Age (X axis) plotted against AUC for IgM anti-S (Y axis) r=0.0565 p=0.4934. **g**, Duration of symptoms in days (X axis) plotted against AUC for IgG anti-RBD (Y axis) r=0.1525, p=0.0633. **h**, Duration of symptoms in days (X axis) plotted against AUC for IgM anti-RBD (Y axis) r=-0.3187,p=<0.0001. i, Duration of symptoms in days (X axis) plotted against AUC for IgG anti-S (Y axis) r=0.0329, p=0.6904. **j**, Duration of symptoms in days (X axis) plotted against AUC for IgM antiS (Y axis) r=0.0824, p=0.3177. **k**, Severity of symptoms (X axis) plotted against AUC for IgG anti-RBD (Y axis) r=0.2679 p=0.0010. **l**, Severity of symptoms (X axis) plotted against AUC for IgM anti-RBD (Y axis) r=-0.1943 p=0.0176. **m**, Severity of symptoms (X axis) plotted against AUC for IgG anti-S (Y axis) r=0.1187 p=0.1492. **n**, Severity of symptoms (X axis) plotted against AUC for IgM anti-S (Y axis) r=0.1597 p=0.0517. All correlations were analyzed by two-tailed Spearman’s.

**Extended Data Figure 4.**
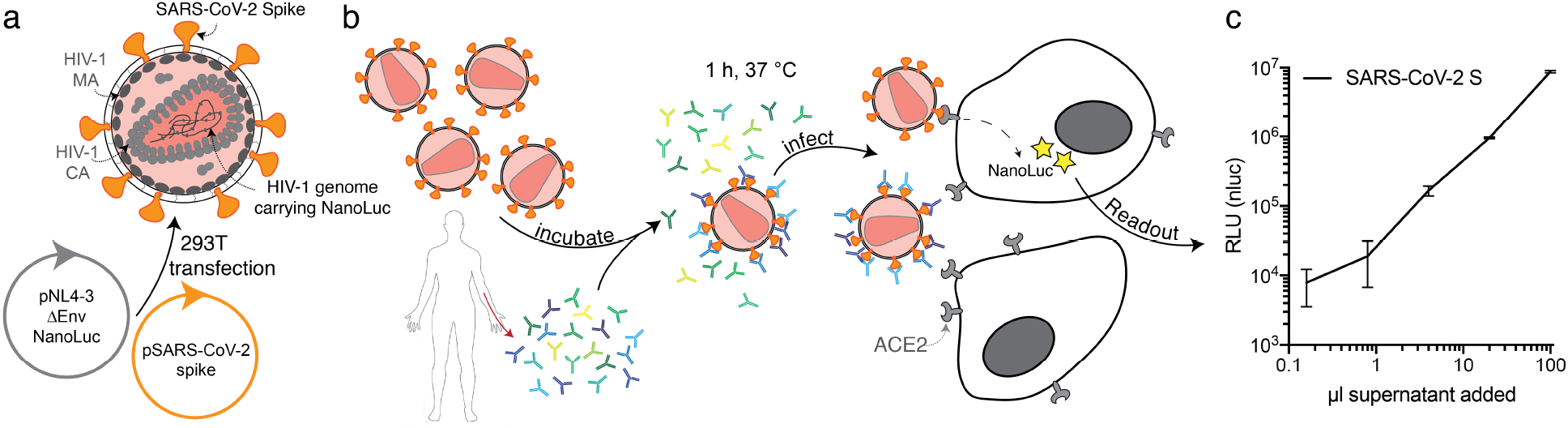
Diagrammatic representation of the SARS-CoV2 pseudovirus luciferase assay. **a**, Co-transfection of pNL4-3ΔEnv-nanoluc and pSARS-CoV-2 spike vectors into 293T cells leads to production of SARS-CoV-2 Spike-pseudotyped HIV-1 particles (SARS-CoV-2 pseudovirus) carrying the *Nanoluc* gene. **b**, SARS-CoV-2 pseudovirus is incubated for 1 h at 37°C with plasma or monoclonal antibody dilutions. The virus-antibody mixture is used to infect ACE2-expressing 293T cells, which will express nanoluc Luciferase upon infection. c, Relative luminescence units (RLU) reads from lysates of ACE2-expressing 293T cells infected with increasing amounts of SARS-CoV-2 pseudovirus.

**Extended Data Figure 5.**
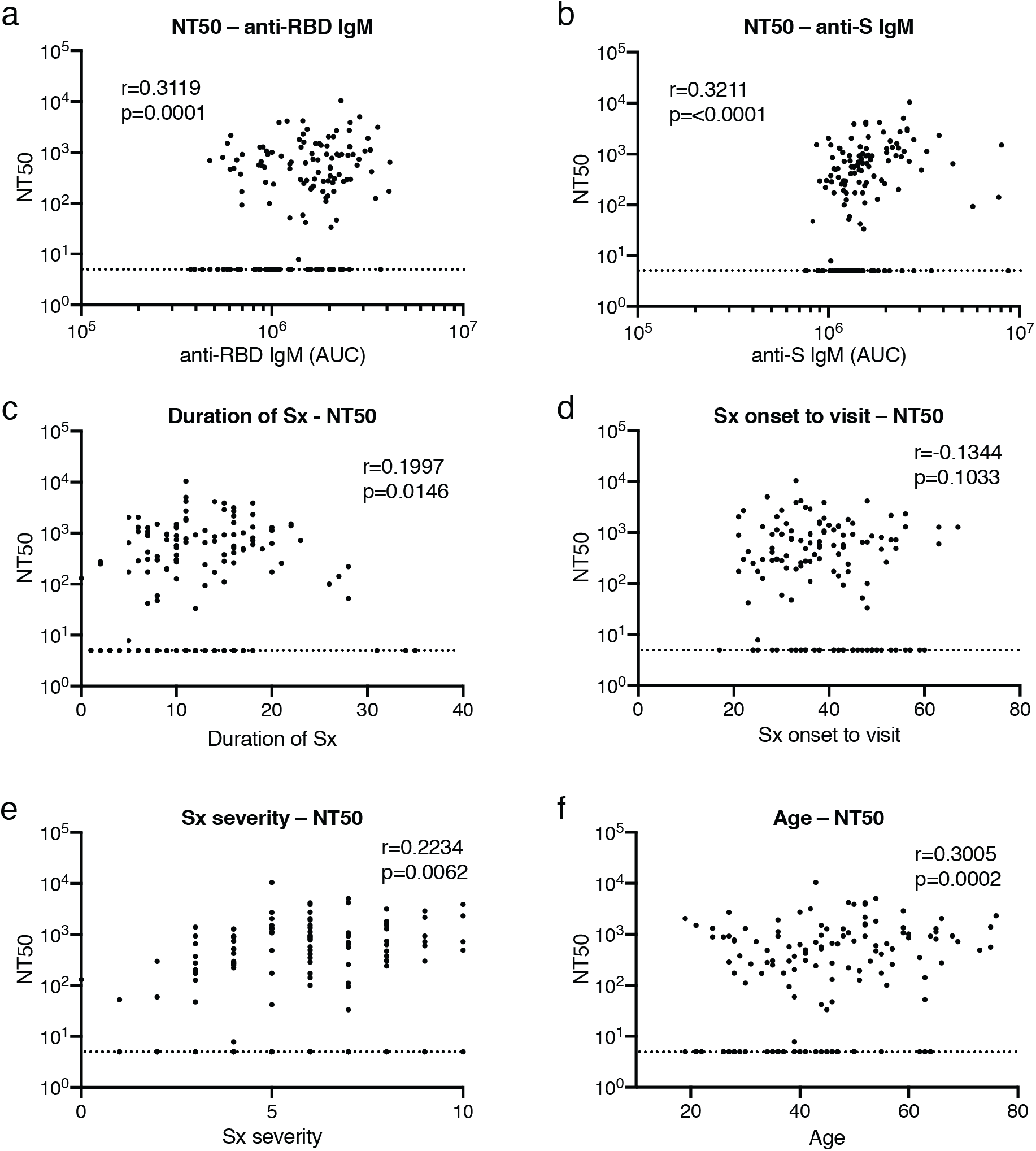
Clinical correlates of neutralization. **a**, Anti-RBD IgM AUC (X axis) plotted against NT_50_ (Y axis) r=0.3119, p=0.0001. **b**, Anti-S IgM AUC (X axis) plotted against NT_50_ (Y axis) r=0.3211, p=<0.0001. **c**, Duration of symptoms in days (X axis) plotted against NT_50_ (Y axis) r=0.1997, p=0.0146. **d**, Time between symptom onset and plasma collection in days (X axis) plotted against NT_50_ (Y axis) r=-0.1344, p=0.1033. **e**, Symptom severity (X axis) plotted against NT_50_ (Y axis) r=0.2234, p=0.0062. **f**, Age (X axis) plotted against NT_50_ (Y axis) r=0.3005, p=0.0002. All correlations were analyzed by two-tailed Spearman’s. Dotted line (NT_50_=5) represents lower limit of detection (LLOD) of pseudovirus neutralization assay. Samples with undetectable neutralizing titers were plotted at LLOD.

**Extended Data Figure 6.**
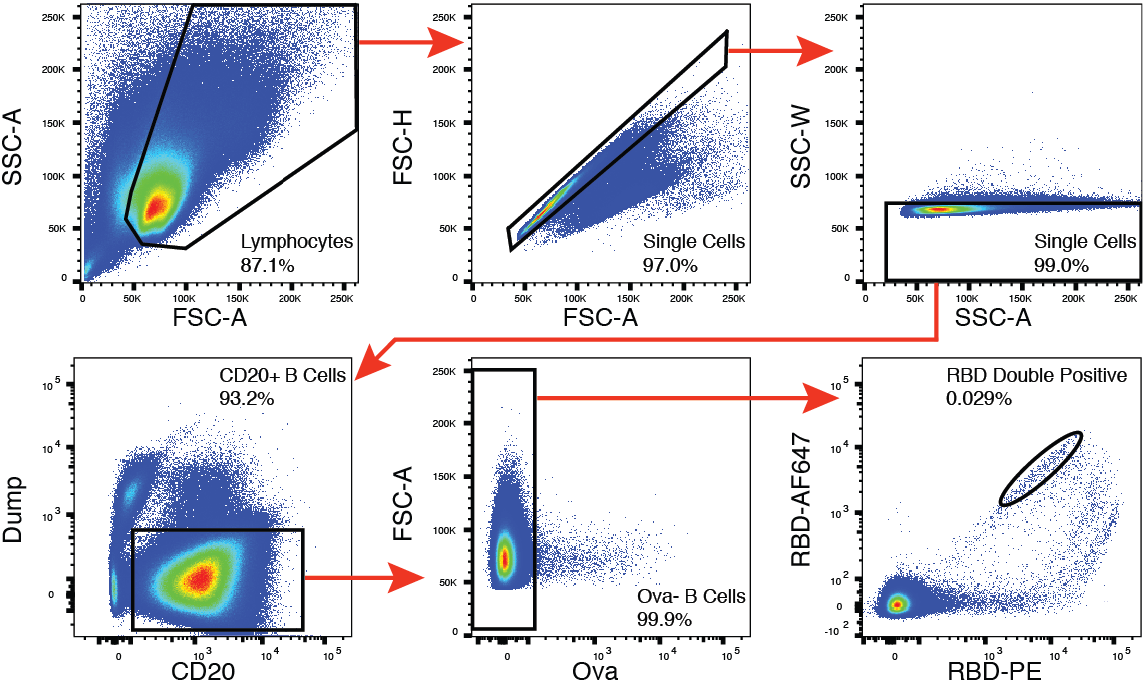
Flow cytometry. Gating strategy used for cell sorting. Gating was on singlets that were CD20^+^ and CD3^-^CD8^-^CD16^-^Ova^-^. Sorted cells were RBD-PE^+^ and RBD-AF647^+^.

**Extended Data Figure 7.**
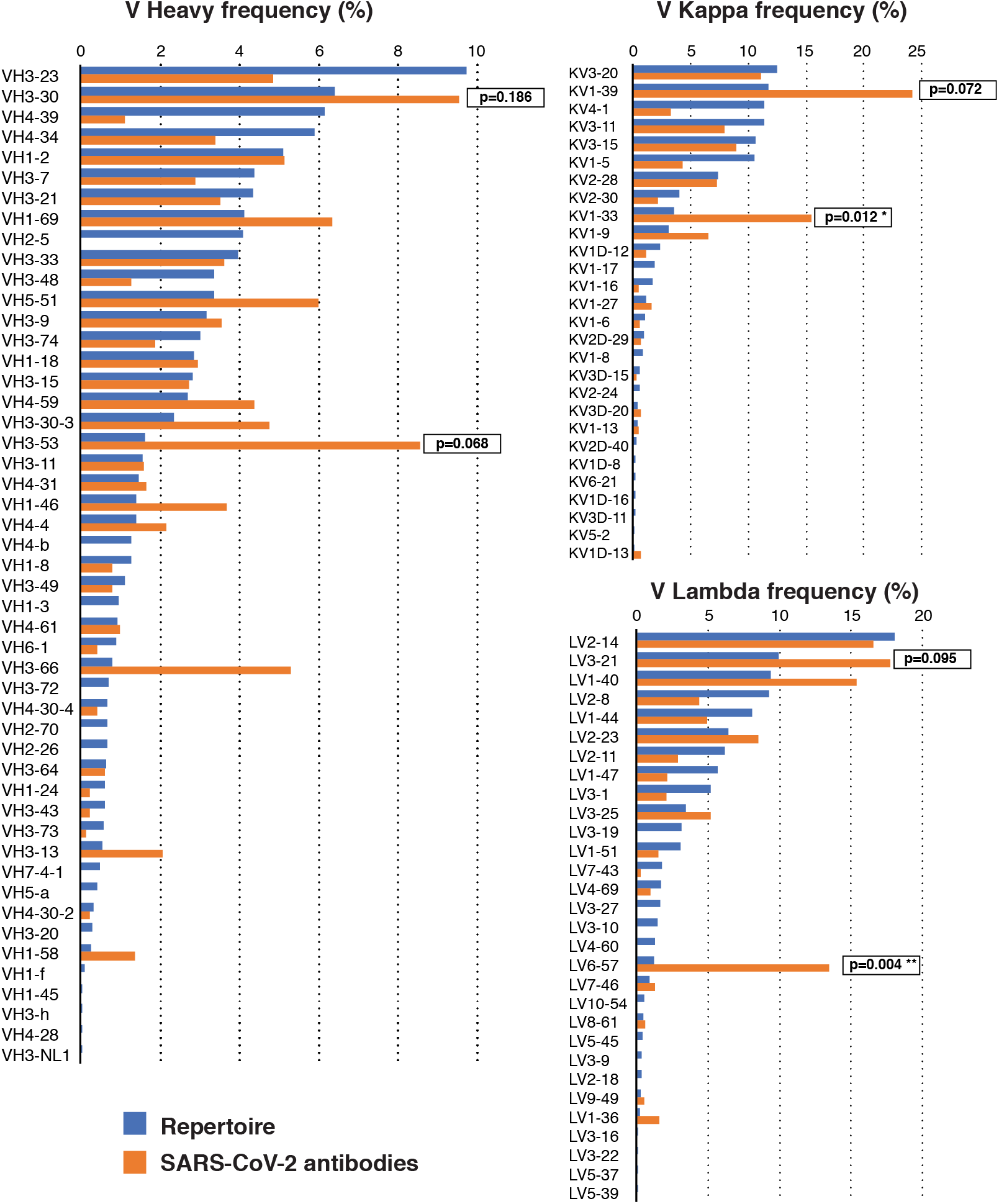
Frequency distributions of human V genes. The two-tailed t test with unequal variance was used to compare the frequency distributions of human V genes of anti-SARS-CoV-2 antibodies from this study to Sequence Read Archive SRP010970^34^.

**Extended Data Figure 8.**
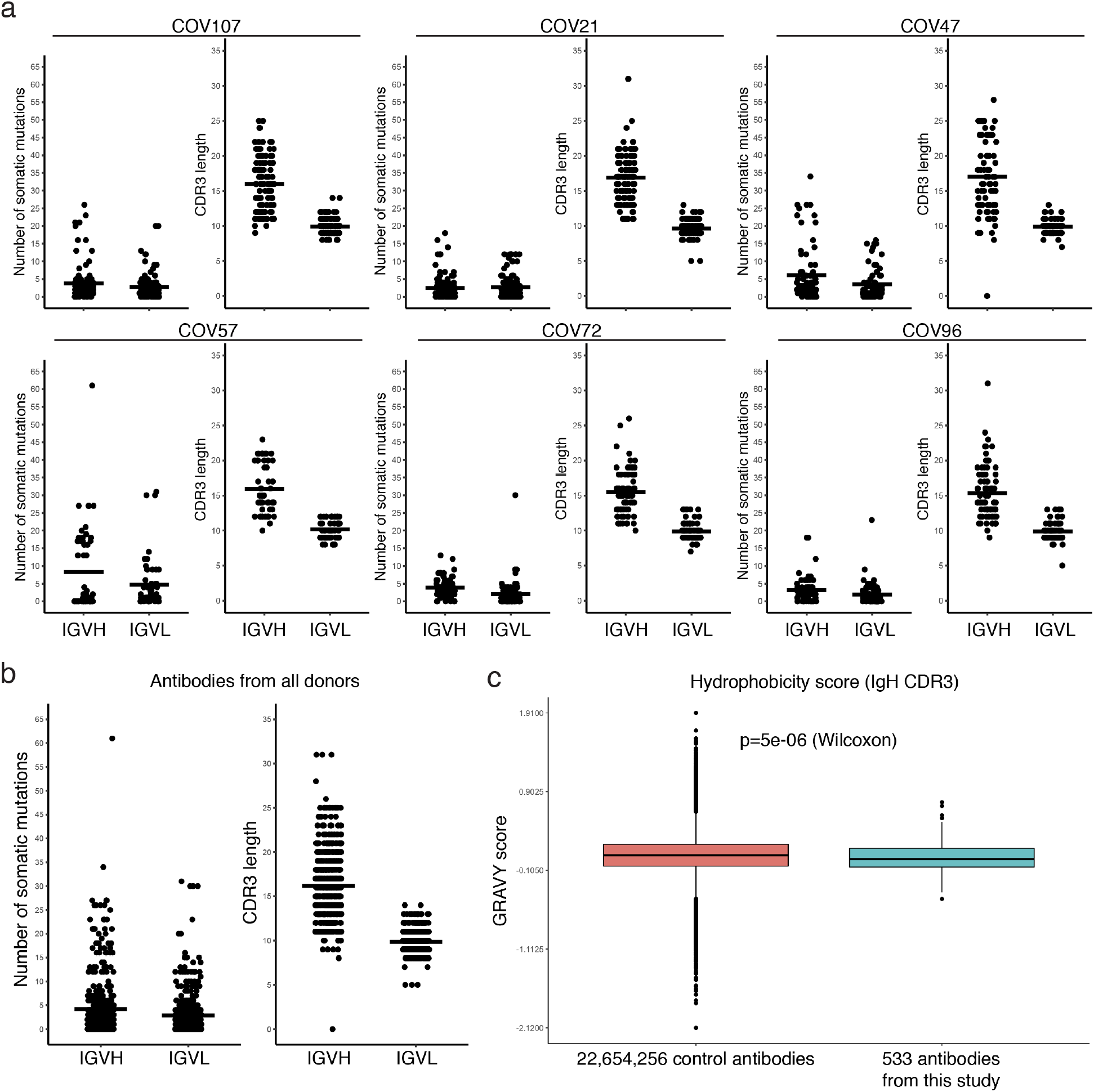
Analysis of antibody somatic hypermutation and CDR3 length. **a,** For each individual, the number of somatic nucleotide mutations (Y axis) at the IGVH and IGVL are shown on the left panel, and the amino acid length of the CDR3s (Y axis) are shown on the right panel. The horizontal bar indicated the mean. **b,** same as in a but for all antibodies combined. **c**, Distribution of the hydrophobicity GRAVY scores at the IGH CDR3 in antibody sequences from this study compared to a public database (see Methods).

**Extended Data Figure 9.**
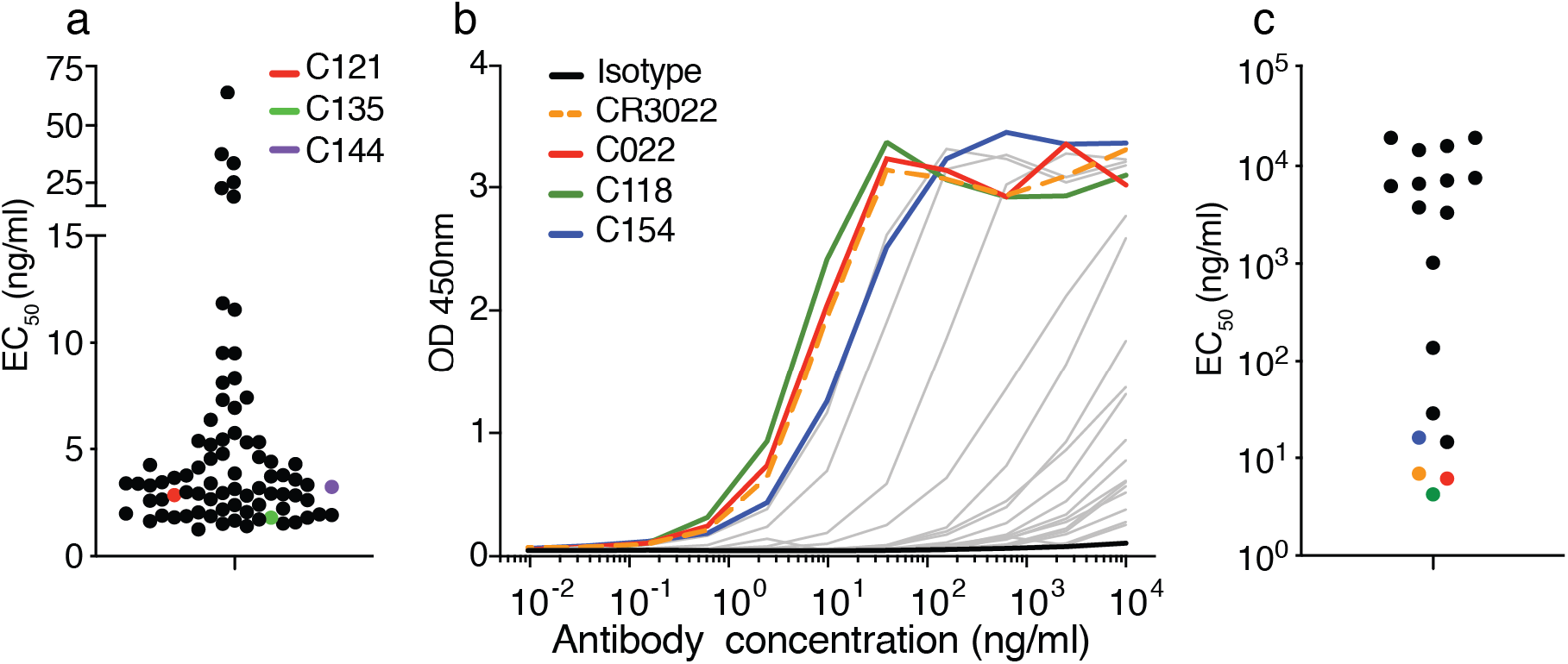
Binding of the monoclonal antibodies to the RBD of SARS-CoV-2 and SARS-CoV. **a**, EC_50_ values for binding to the RBD of SARS-CoV-2. **b and c**, Binding curves and EC_50_ values for binding to the RBD of SARS-CoV.

**Extended Data Figure 10.**
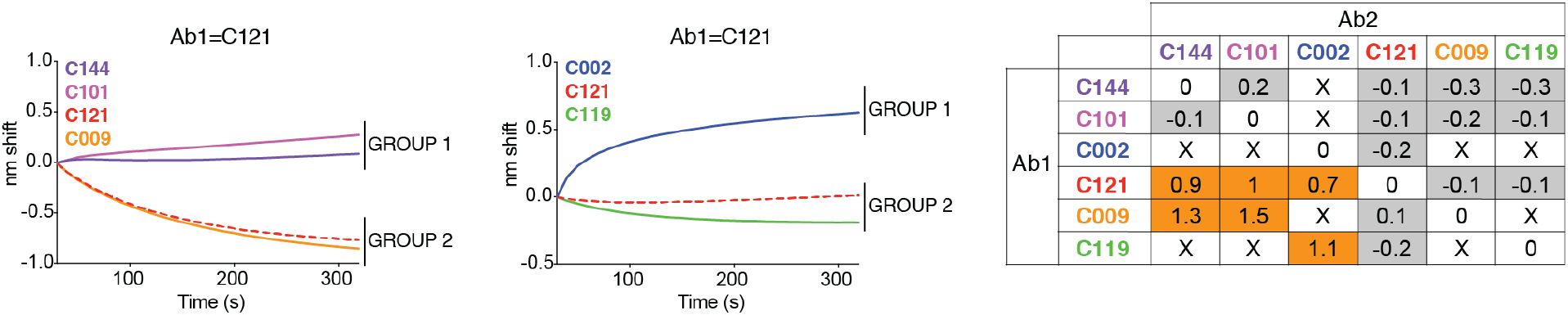
Biolayer interferometry experiment. showing binding of antibodies C144, C101, C002, C121, C009, C019 (see also main text Fig. 4). Graphs show secondary antibody binding to preformed C121 IgG-RBD complexes. The table displays the shift in nanometers after second antibody (Ab2) binding to the antigen in the presence of the first antibody (Ab1). Values are normalized by the subtraction of the autologous antibody control.

